# A bright synthetic near-infrared luciferin enhances the capabilities of deep-tissue bioluminescence imaging using firefly luciferases

**DOI:** 10.1101/2025.01.20.633992

**Authors:** Ryohei Saito-Moriya, Satoshi Iwano, Nobuo Kitada, Norihisa Yamasaki, Genta Kamiya, Katsunori Ogo, Takashi Sugiyama, Ariunbold Chuluun-Erdene, Takayuki Isagawa, Rika Obata, Ryosuke Ijuin, Hideki Abe, Yuji Kashiwakura, Yo Mabuchi, Megumu Takahashi, Yoshihiro Yamamoto, Daigo Sawaki, Tatsuyuki Sato, Shigeru Sato, Hidekazu Nishikii, Kotaro Yoshimura, Hiroyuki Hioki, Tsukasa Ohmori, Toshiaki Nakashiba, Takashi Hirano, Hiroshi Aoyama, Norihiko Takeda, Atsushi Miyawaki, Shojiro A. Maki, Takahiro Kuchimaru

## Abstract

Synthetic bioluminescence reactions exhibiting near-infrared (NIR)-shifted spectra have been explored to improve deep-tissue imaging through the design of firefly luciferin analogues. Although the NIR bioluminescence reactions improve the tissue penetration of bioluminescence signals from deep tissues, their photon output is markedly lower compared to the natural reaction with D-luciferin and firefly luciferase (Fluc), often by an order of magnitude or more. Consequently, in most instances, the sensitivity of NIR bioluminescence imaging (NIR-BLI) has not yet substantially surpassed that of BLI with the natural firefly reaction. Here, we present a synthetic firefly luciferin, named AkaSuke, that generates intense NIR bioluminescence (λ_max_ = 680 nm) in reaction with Fluc, greatly improving the detection sensitivity beyond that of the D-luciferin/Fluc reaction for targeting deep tissue. AkaSuke enables sensitive visualizations of ectopic hematogenesis through entire tissues of mice over time following transplantation of bone marrow stem cells labeled with Fluc. We additionally identify a Japanese firefly luciferase, DkumLuc1, that displays higher catalytic activities for bioluminescence emission of AkaSuke compared to typical Fluc, resulting in detection sensitivity comparable to that of AkaLumine/Akaluc reaction, one of the most sensitive bioluminescence systems for deep tissue imaging. We further propose the potential of the AkaSuke/DkumLuc1 reaction as an orthogonal pair with the AkaLumine/Akaluc for sensitive dual-target tracking in mice. Overall results suggest that AkaSuke enhances the capabilities of deep-tissue bioluminescence imaging using Fluc and its variant, and could serve as an emerging benchmark for the molecular design of NIR luciferin analogues.

## Introduction

Bioluminescence imaging (BLI) is employed in broad range of biomedical studies with small animals because of its non-invasiveness and high sensitivity ^1–3^. To maximize BLI performance in animals, a suitable luciferin/luciferase pair has been explored. Among several luciferin-luciferase pairs available in scientific researches, the pair of D-luciferin and firefly luciferase (Fluc) has been a gold standard for non-invasive imaging in small animals. This is primarily because D-luciferin is highly biocompatible with its high water solubility, stability in blood circulation and relatively ubiquitous diffusion in animal tissues ^4^. Consequently, major efforts have been directed toward extending the visible bioluminescence wavelength of D-luciferin (λ_max_ = 560 nm) to near-infrared (NIR) region (>650 nm), where light penetration into biological tissues is more efficiently, thereby enhancing deep tissue imaging capabilities ^5,6^. We and others have successfully identified firefly luciferin analogues that generate NIR bioluminescence in the reaction with Fluc ^7–11^. However, current NIR luciferin analogues significantly lacked photon output compared to D-luciferin in reaction with Fluc, which does not substantially improve detection sensitivity of BLI using the D-luciferin/Fluc reaction. To address this issue, we engineered Fluc to create Akaluc that harnesses highly improved photon output with a NIR luciferin analogue, AkaLumine ^12^. The AkaLumine/Akaluc BLI exceptionally surpasses detection sensitivity of the D-luciferin/Fluc BLI for deep-tissue imaging.

Another critical demand in NIR-BLI is the development of an orthogonal bioluminescence reaction for visualizing dual or more targets in living tissues. While only a few dual NIR-BLI systems have been successfully developed, the exploration of orthogonal pairs of firefly luciferin analogues and mutant luciferases remains an active area of research ^13,14^. Even in the feasible dual NIR-BLI system ^15^, the bottle neck is the lack of photon output from NIR bioluminescence reactions, which impedes highly sensitive BLI in animals. Therefore, there is a need for further development of dual NIR-BLI systems to enhance the capabilities of deep-tissue BLI using firefly luciferases.

In this study, we synthesized a firefly luciferin analogue, named AkaSuke, by introducing thiazole ring in AkaLumine. AkaSuke displayed robust wavelength shift in NIR wavelength region (λ_max_ = 680 nm) in reaction with Fluc. Notably, the AkaSuke/Fluc reaction generated photons efficiently that is almost with an order of magnitude higher that the AkaLumine/Fluc reaction in living cells. As a result, AkaSuke successfully detected cancer cells and hematopoietic cells in deep tissues with several-fold higher sensitivity of D-luciferin. We further identified DkumLuc1, one of the Japanese firefly luciferases, that highly catalyzes photon emission of AkaSuke, resulting in comparable imaging sensitivity with AkaLumine/Akaluc reactions in deep tissue imaging. Interestingly, DkumLuc1 scarcely catalyzes AkaLumine. Thus, the potential of an orthogonal pairs of AkaSuke/DkumLuc1 and AkaLumine/Akaluc is demonstrated by tracking cancer cells and T cells in mice.

## Results

### Design of AkaSuke and its analogues

We designed AkaSuke (**1a**) as a bioisostere compound by fusing the thiazole ring structure of D-luciferin with the butadienyl moiety of AkaLumine. This design is crucial for enhancing bioluminescence photon output and for extending the wavelength of bioluminescence (Fig. 1a). In parallel, we designed AkaSuke-like analogues **1b** and **1c** to gain insights on the structure-activity relationships of luciferin/luciferase reaction (Fig. 1a). In order to verify whether AkaSuke (**1a**) and its analogues **1b** and **1c** designed with a dimethylamino-substituted 5-membered ring (thiazole and thiadiazole) as the main fluorophore moiety also have sufficient luminescence properties, the luminescence properties of **1a–c** were predicted by density functional theory (DFT) and time-dependent DFT (TD-DFT) calculations of the oxyluciferin forms (oxy-**1a–c**) corresponding to **1a–c** (Supplementary Fig. 1). The HOMO-LUMO energy gaps (Δ*E*_H-L_) and other parameters (the wavelengths (λ_tr_), oscillator strengths (*f*), and configurations for the S_0_→S_1_ transitions) for oxy-**1a–c** were resembled those of the previously reported NIR luciferin analogues such as AkaLumine and seMpai (Fig. 1a) ^10,16^, and their *f* values of ca.0.7 would predict that the excited oxy-**1a–c** have sufficient fluorescence ability in the bioluminescence reactions (Supplementary Fig. 1 and Supplementary Table 1-5). In addition, the λ_tr_ value of AkaSuke (**1a**) was anticipated to be extended to longer wavelengths compared to AkaLumine and seMpai. Then, AkaSuke (**1a**) and its analogues **1b** and **1c** were synthesized by 7-step reactions, respectively (Supplementary Fig. 2).

**Fig. 1.**
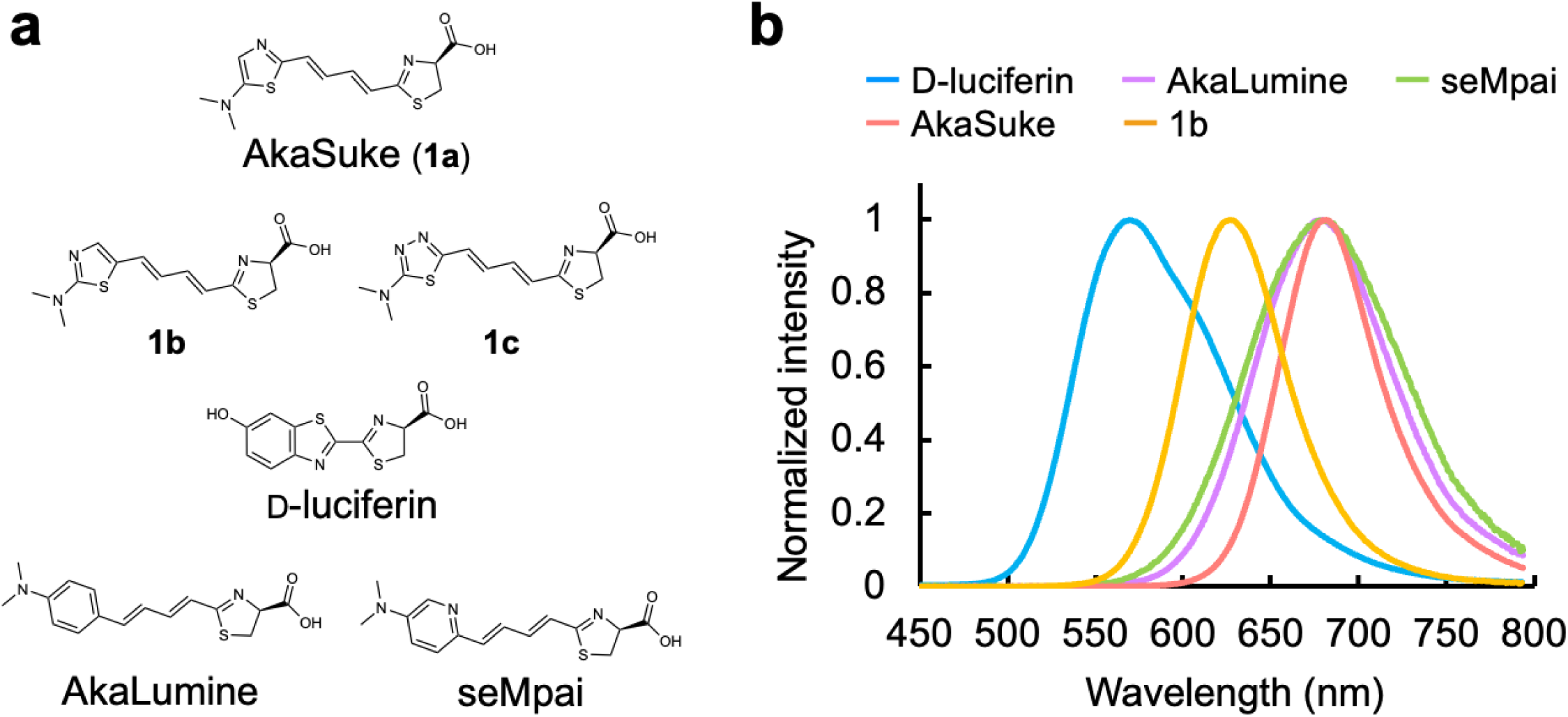
Synthetic firefly luciferin analogues with an extended π conjugation and a 5-membered ring. **a** Molecular structure of the synthetic luciferin AkaSuke, **1b** and **1c**. The structure of D-luciferin, AkaLumine and seMpai is also shown. **b** Bioluminescence spectrum of the substrates in reaction with natural Fluc. Source data are provided as a Source Data file.

Bioluminescence properties of AkaSuke and its analogues were characterized with the North American firefly luciferase (*Photinus pyralis*) (Fluc). Notably, the bioluminescence properties of AkaSuke were much unique when compared to its analogues (Table 1). In reaction with Fluc, the photon emission of AkaSuke was approximately 3.4 folds higher than that of AkaLumine, while it was approximately 0.7-folds weaker than that of D-luciferin (Table 1). The bioluminescence spectra of AkaSuke, when reacted with Fluc, displayed emission maximum at 680 nm (Fig. 1b and Table 1). The emission peak of AkaSuke is notably red-shifted in comparison to the peak of D-luciferin. AkaSuke’s emission maximum is comparable to that of AkaLumine and seMpai. Unlike the D-luciferin/Fluc reaction, the NIR bioluminescence spectrum of the AkaSuke/Fluc reaction is stable across the varied pH conditions (Supplementary Fig.3). The *K*_m_ and *V*_max_ values of AkaSuke in reaction with Fluc was 118 μM and 1.40 ×10^7^, respectively (Table 1). These values are closely aligned with those of D-luciferin rather than those of AkaLumine, indicating that AkaSuke inherits favorable photon output properties of D-luciferin with NIR-shifted wavelength. In contrast, the analogue **1b** generated efficient photons, while modest red-shift in wavelength (λ = 620 nm) compared to AkaSuke (Fig. 1b and Table 1). The analogue **1c** failed to produce detectable bioluminescence (Table 1). Additionally, UV-visible spectrophotometry analysis revealed that AkaSuke exhibits exceptional solubility, reaching concentrations up to 545 mM in phosphate-buffered saline (PBS, pH = 7.4) (Supplementary Fig. 4). This solubility is much higher than that of the AkaLumine analogue, which exbibits a solubility of 30 mM in neutral pH buffer^10^. Moreover, it exceeded the typical solubility range (100 – 300 mM) observed for D-luciferin salts^17,18^. Collectively, these results indicated that AkaSuke holds significant promise as a potent NIR luciferin analogue for BLI applications in biomedical studies.

**Table 1.**
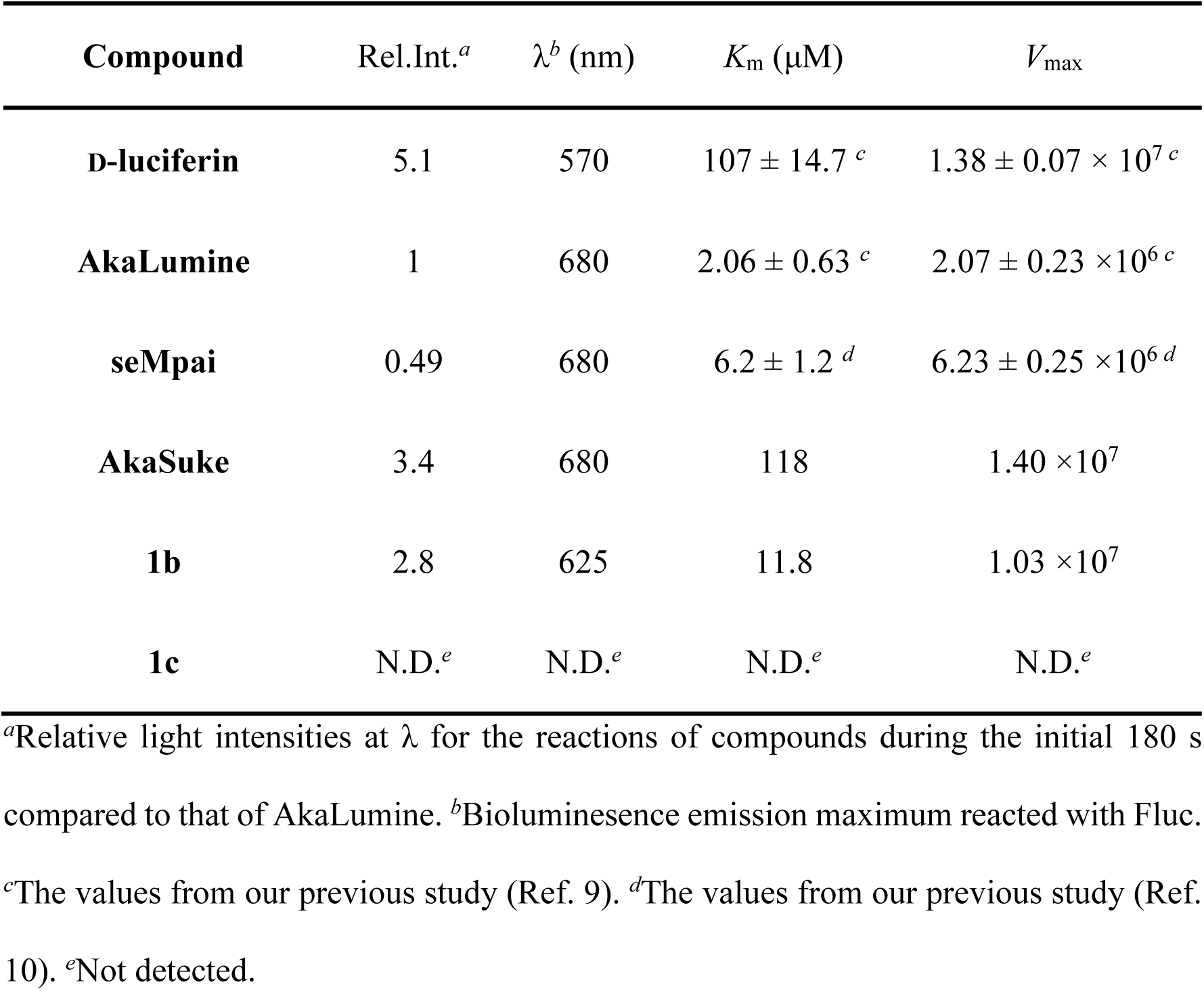
Bioluminescence properties of D-luciferin and the luciferin analogues in the reaction with Fluc.

### Characterization of AkaSuke in mammalian cells

We tested whether AkaSuke is able to efficiently produce bioluminescence in living cells. To do so, we established murine breast cancer E0771/Venus-Fluc cells that constitutively express yellow fluorescent protein (Venus) fused with Fluc (luc2 standard Fluc variant for mammalian cells). We treated E0771/Venus-Fluc with increasing concentrations of AkaSuke, D-luciferin and AkaLumine-HCl. In consistent with the purified Fluc protein assay (Table 1), AkaSuke generated intense bioluminescence photons which are 8-fold higher than AkaLumine-HCl (Fig. 2a). The bioluminescent signals in NIR wavelength range, AkaSuke generated much higher bioluminescent signals than D-luciferin and AkaLumine-HCl (Fig. 2b). Consistent with these results, AkaSuke outperformed seMpai that is an AkaLumine analogue showing higher solubility in neutral pH buffer^10^ (Supplementary Fig 5). As similar with current luciferin-luciferase pairs, AkaSuke linearly generated photon output with increased number of cells expressing Fluc (Fig. 2c). While, unlike AkaLumine-HCl, the bioluminescence kinetics of AkaSuke exhibited a relatively slow reaction rate, yet it was faster than that of D-luciferin at unsaturated substrate concentrations (Fig. 2d). Taken together, AkaSuke significantly improved NIR bioluminescence outputs over D-luciferin and AkaLumine-HCl in living cells expressing Fluc.

**Fig. 2.**
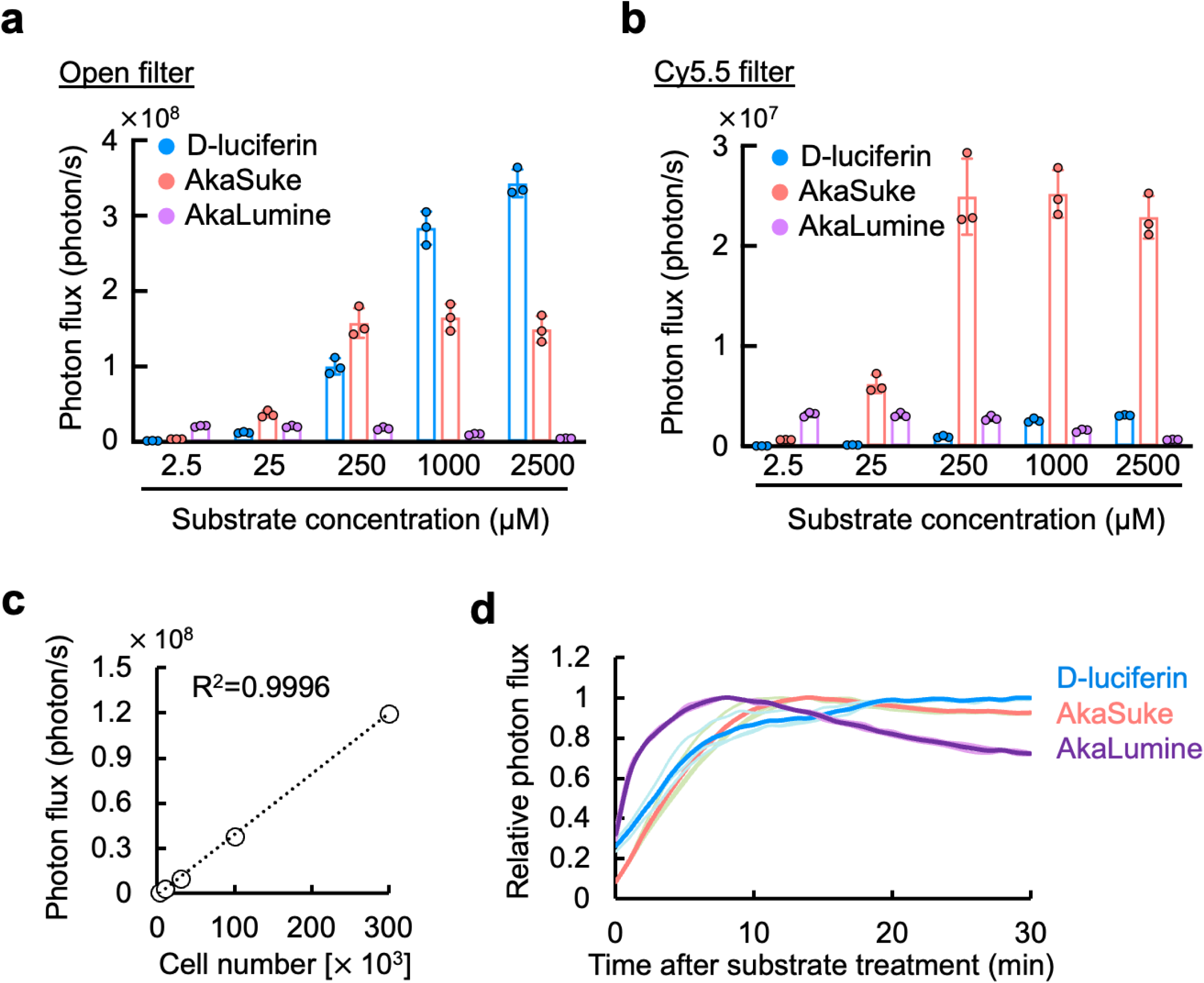
Characterization of AkaSuke in living cells. **a** Bioluminescence emission from living cells expressing Fluc (E0771/Venus-Fluc) in reaction with the substrate at indicated concentrations. Bioluminescence emissions were measured using an open filter. Data were statistically analyzed with two-tailed Student’s t-test (n = 3 biologically independent samples). Data are presented as mean values ± SEM. Source data are provided as a Source Data file. **b** Bioluminescence emission from E0771/Venus-Fluc was measured using a near-infrared (Cy5.5) filter. Data were statistically analyzed with two-tailed Student’s t-test (n = 3 biologically independent samples). Data are presented as mean values ± SEM. Source data are provided as a Source Data file. **c** The correlation between Fluc-expressing cell number and bioluminescence emission of AkaSuke. E0771/Venus-Fluc cells were treated with 250 μM AkaSuke. Data are presented as mean values (n = 3 biologically independent samples). Source data are provided as a Source Data file. **d** The bioluminescence kinetics of substrates in living cells expressing Fluc. E0771/Venus-Fluc cells were treated with the substrate concentration of 25 μM for D-luciferin and AkaSuke, 1 μM for AkaLumine. Thick lines represent the mean values, which are calculated from individual thin lines (n = 3 independent biological samples). Source data are provided as a Source Data file. All experiments were repeated at least twice, and similar results were reproducible.

### Animal imaging with AkaSuke

We then sought to assess performance of AkaSuke for deep tissue imaging in animals. We firstly determined suitable intraperitoneal administration dose of AkaSuke for animal imaging. BLI with varied administration dose indicated that the dose of 3 μmol/body (equivalent to 46.5 mg/kg for a 20 g body weight) saturated bioluminescence outputs in Fluc-expressing cells trapped in lung capillaries of the mice (Supplementary Fig. 6a and b). When administered to a CAG-luc2 knockin mouse at the ROSA26 locus in which Luc2 is expressed throughout the body (ROSA CAG-luc2 KI mouse), the administration of 3 μmol/body of AkaSuke yielded more intense NIR bioluminescence than the case of AkaLumine at the same dose (Supplementary Movies 1 and 2). The signal intensity was strong enough to be recorded by a consumer-grade digital color camera in far-red color, facilitating real-time color BLI of freely-behaving mice. Next, we prepared mice bearing overt lung metastasis by intravenous injection of E0771/Venus-Fluc cells to quantitively evaluate the detection sensitivity of AkaSuke over D-luciferin that has been current standard substrate for BLI of animals. The comparison of the substrates was performed with bioluminescence images of same mice bearing E0771/Venus-luc2 lung metastasis after intraperitoneal injection in order of D-luciferin and AkaSuke at a 6-hour interval. Notably, AkaSuke greatly enhanced the signal from the lung metastases, as demonstrated by 7.7-fold higher with AkaSuke compared with standard 10 μmol dose of D-luciferin (Fig. 3a and b). AkaSuke displayed fast peak of bioluminescence emission (Supplementary Fig. 6c). While, its penetration into blood brain barrier (BBB) is moderate relative to AkaLumine, AkaSuke still holds superior sensitivity over D-luciferin in deep brain tissue imaging targeting striatum with the AAV2/1-SynTetOff-Venus-Fluc construct ^19^ (Supplementary Fig. 7). Additionally, we observed that AkaSuke induced lower levels of autoluminescence in the liver compared to AkaLumine-HCl (Supplementary Fig. 8a and b). Concomitantly, AkaSuke failed to enhance bioluminescence emission in liver parenchyma (Supplementary Fig. 8c and d); however, it enabled sensitive detection of liver metastasis in mice (Supplementary Fig. 8e and f). These observations motivated us to test detection sensitivities of AkaSuke in multiple organs in animal models. We constructed a hematopoietic expansion model by transplantation of bone marrow cells (BM cells) harvested from transgenic mice ubiquitously expressing a reporter gene of Fluc fused with cpVenus (*ff*luc) ^20^. Bone marrow cells of *ff*luc mice contained c-Kit+ Sca-1+ hematopoietic stem cells that robustly express the reporter gene (Supplementary Fig. 9a). After transplantation of BM cells via tail vein of immune deficient mice, repeatedly injection of AkaSuke successfully visualized hematopoietic expansion through the entire body of mice (Fig. 3c). In consistent with cancer metastasis model, AkaSuke improved detection sensitivities of hematopoietic cell accumulation in lymph nodes and hind limb bone marrow over D-luciferin (Supplementary Fig. 9b and Fig. 3d and e). These results demonstrate that AkaSuke overall improves sensitivity of targets in multiple deep organs over D-luciferin.

**Fig. 3.**
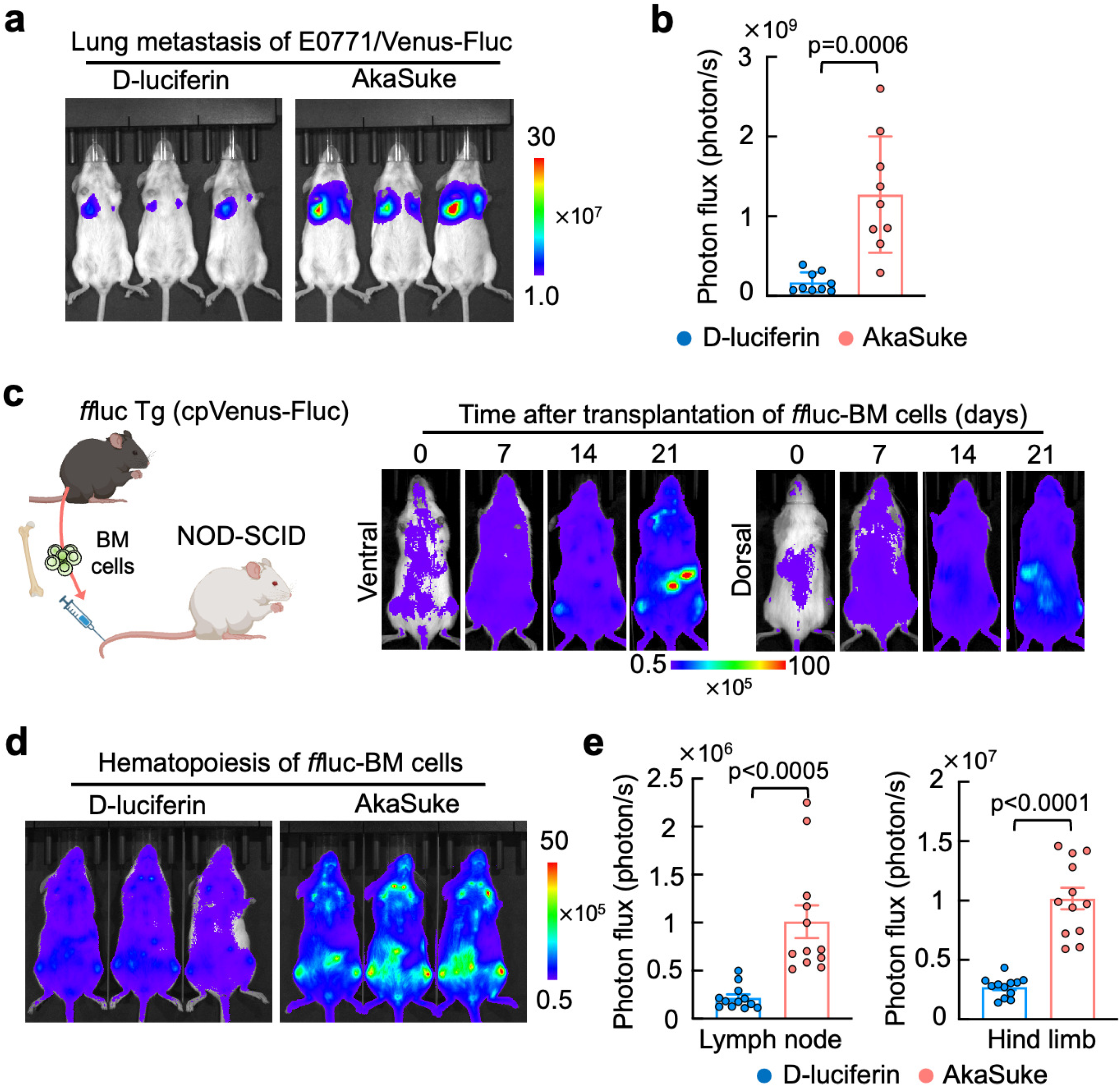
NIR-BLI of living mice using the AkaSuke/Fluc rection. **a** Representative bioluminescence images of mice harboring E0771/Venus-Fluc lung metastasis obtained by D-luciferin and AkaSuke in same mice. The mice were intraperitoneally injected with D-luciferin (10 μmol/body) and AkaSuke (3 μmol/body). **b** Quantitative analysis of bioluminescence emission from lung metastasis generated with D-luciferin and AkaSuke (n = 8 independent metastatic lesions in 5 mice, p-value for D-luciferin vs AkaSuke, two-tailed Student’s *t* test). Source data are provided as a Source Data file. **c** BLI of hematopoietic expansion process using AkaSuke. The experimental diagram (left) and representative time-course bioluminescence images obtained by repeatedly injecting AkaSuke (3 μmol/body) (right panel). Bone marrow cells (BM cells) were isolated from transgenic mice expressing Fluc (*ff*luc Tg mice) and transplanted into wild-type NOD-SCID mice. **d** Representative bioluminescence images of hematopoiesis obtained by D-luciferin and AkaSuke in same mice. The mice were intraperitoneally injected with i.p. injection of D-luciferin (10 μmol/body) and AkaSuke (3 μmol/body) after transplanting BM cells isolated from *ff*luc Tg mice. **e** Quantitative analysis of bioluminescence emission from lymph nodes (LN) and hind limbs (HL) obtained with D-luciferin and AkaSuke (n = 12 independent LN and HL in 6 mice, p-value for D-luciferin vs AkaSuke, two-tailed Student’s *t* test). Source data are provided as a Source Data file. All experiments were repeated at least twice, and similar results were reproducible.

### Enhanced animal imaging with AkaSuke and firefly luciferase variant

We found that the *Drilaster* luciferase derived from a firefly in south islands of Japan, named DkumLuc1^21^, displayed higher catalytic activity of D-luciferin than that of Fluc in living cells (Supplementary Fig. 10), and robust NIR shift in reaction with AkaSuke (Fig. 4a). As similar with the AkaSuke/Fluc reactions, the bioluminescent spectrum emitted by AkaSuke/DkumLuc1 reactions are less sensitive to varied pH conditions (Supplementary Fig. 11). This motivated us to compare the BLI performance of the AkaSuke/DkumLuc1 with that of the AkaLumine/Akaluc that is currently one of the best bioluminescence reactions for animal imaging. In living cells, the AkaSuke/DkumLuc1 reaction yielded bioluminescence outputs comparable to that of the AkaLumine/Akaluc reaction (Fig. 4b). Consistently, AkaSuke/DkumLuc1 reaction demonstrated similar detection sensitivity to the AkaLumine/Akaluc reaction for cells disseminated in the lung (Fig. 4c and d), indicating that the AkaSuke/DkumLuc1 system holds the capability for highly sensitive modality suitable for deep-tissue imaging.

**Fig. 4.**
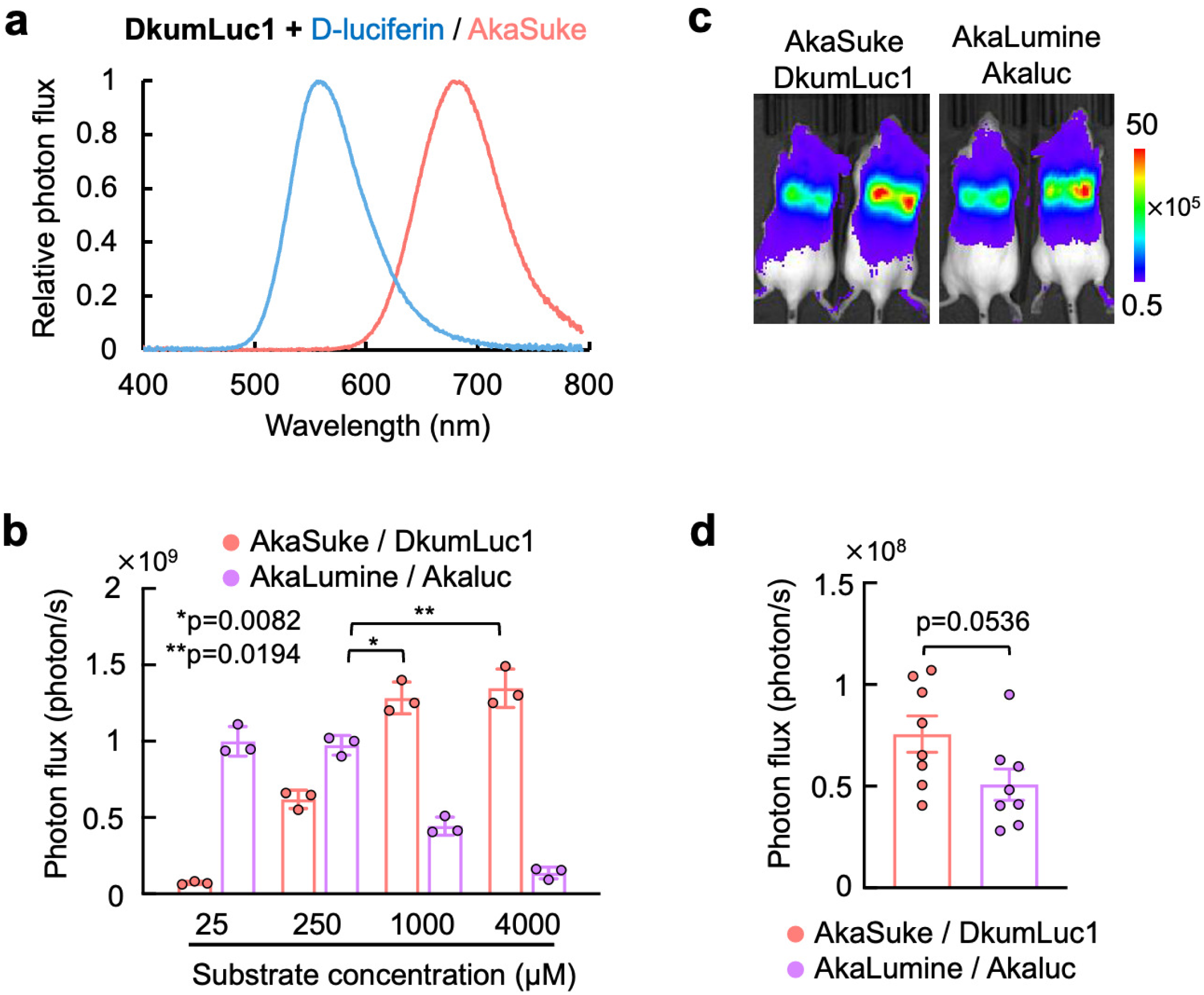
Enhanced deep tissue BLI using AkaSuke/DkumLuc1. **a** Bioluminescence spectrum of DkumLuc1 in reaction with D-luciferin and AkaSuke. **b** Bioluminescence emission from living HEK293T cells transiently expressing DkumLuc1 or Akaluc treated with AkaSuke or AkaLumine at indicated concentrations (n = 3, p-value for 1000 or 4000 μM AkaSuke vs 250 μM AkaLumine, one-way ANOVA followed by Bonferroni’s multiple comparison). **c** Representative BL images of cells trapped in the lung obtained with the AkaSuke/DkumLuc1 or AkaLumine/Akaluc reaction. HEK293T cells expressing DkumLuc1 or Akaluc were intravenously injected from the tail vein 15 minutes before i.p. administration of AkaSuke (3 μmol/body) or AkaLumine (1.5 μmol/body). **d** Quantitative analysis of BL emission from the lung using the AkaSuke/DkumLuc1 or AkaLumine/Akaluc reaction (n = 8 independent lung tissue in 4 mice, p-value, two-tailed Student’s *t* test).

### Dual-target detection with NIR-BLI

In addition to enhanced catalytic activity for AkaSuke, we found that DkumLuc1 displayed unique properties of bioluminescence reactions with AkaLumine. At the peak intensity under varying substrate concentration, the AkaSuke/DkumLuc1 reaction produced 200-fold higher bioluminescence signals compared to the AkaLumine/DkumLuc1 reaction (Fig. 5a and b). Whereas, the AkaSuke/AkaLuc reaction generated 15-fold lower bioluminescence signals over the AkaLumine/AkaLuc rection, suggesting potential orthogonal capabilities of these bioluminescence reactions for dual-target detection using NIR-BLI (Supplementary Table 6).

**Fig. 5.**
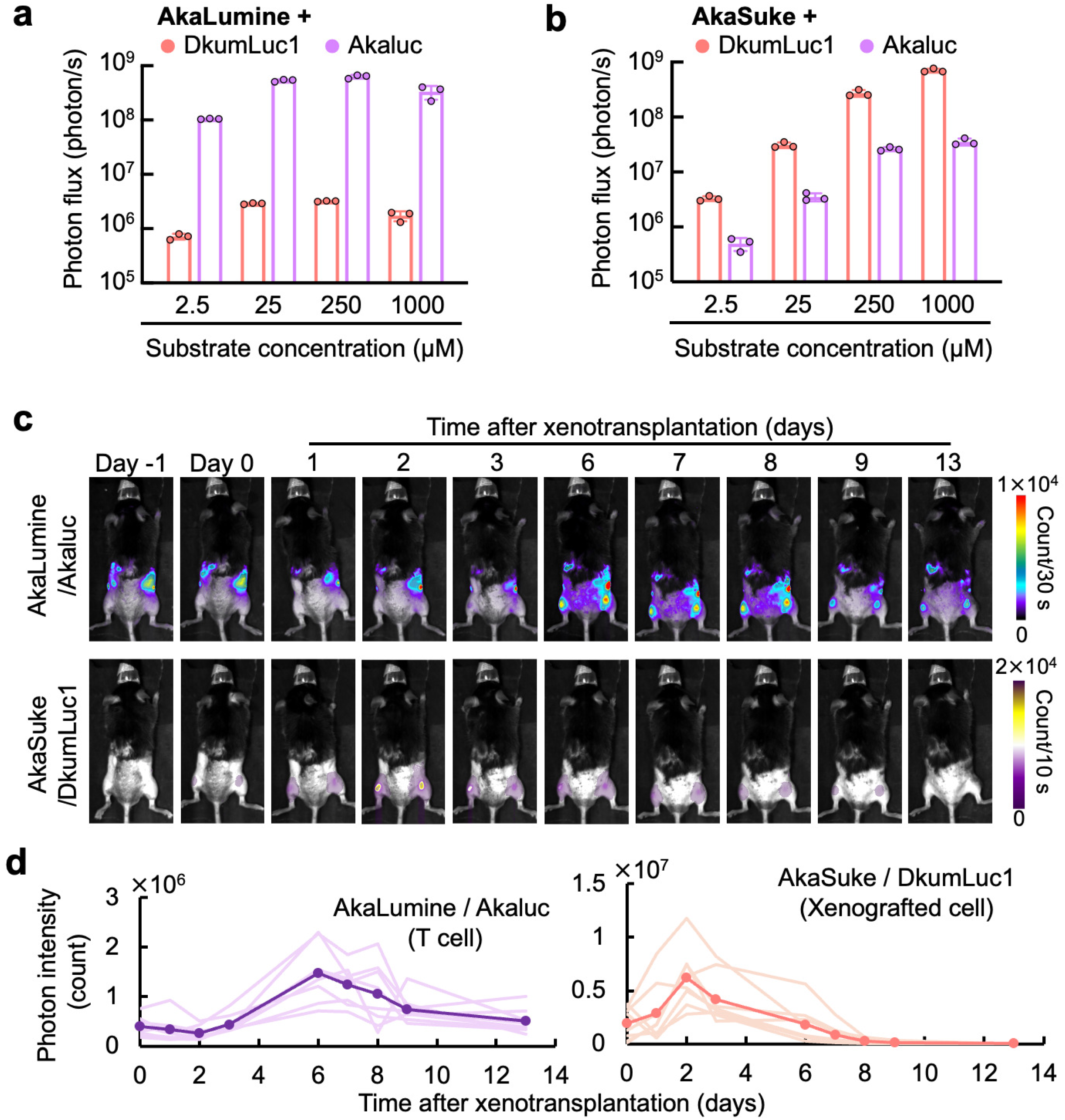
Dual target detection with NIR-BLI by combining AkaSuke/DkumLuc1 and AkaLumine/AkaLuc. **a** Bioluminescence emission from living HEK293T cells expressing DkumLuc1 or Akaluc treated with AkaLumine at indicated concentrations. **b** Bioluminescence emission from living HEK293T cells expressing DkumLuc1 or Akaluc treated with AkaSuke at indicated concentrations. **c** Representative dual NIR-bioluminescence images of T cell migration (AkaLumine/Akaluc) and xenogeneic transplanted cell proliferation (AkaSuke/DkumLuc1). The bioluminescence images for xenotransplant proliferation were acquired by i.p. injection of AkaSuke (3 μmol/body) 6 hours after i.p. injection of AkaLumine (1.5 μmol/body). **d** NIR bioluminescence monitoring of T cell accumulation at xenotransplantation sites (left panel) and xenogeneic grafted cells (right panel). Thick lines represent the mean values, which are calculated from individual thin lines (n = 8 independent xenotransplantation sites in 4 mice). Source data are provided as a Source Data file. All experiments were repeated at least twice, and similar results were reproducible.

To assess the potential of the dual NIR-BLI system, we combined the AkaSuke/DkumLuc1 reaction with Lck-Akaluc mice where T cells are specifically labeled with Akaluc ^22^. Given that AkaSuke exhibited weak but not negligible bioluminescence reactivity with Akaluc in living cells, we investigated the cross reactivity of AkaSuke with Akaluc in Lck-Akaluc mice. In consistent with the cellular assay, AkaSuke elicited weak bioluminescence in Lck-Akaluc mice; however, its contribution was limited to increasing the baseline by no more than 10% (Supplementary Fig. 12). We tested whether the dual NIR-BLI is capable to visualize immune rejection process of the human cells in mice (Supplementary Fig. 13 and Fig. 5c). In consistent with a previous study using positron emission tomography imaging ^23^, the dual NIR-BLI detected persistent T cell recruitment at a xenograft site for a week following xenotransplantation, even subsequence to the initiation of the immune rejection process at an earlier time point (Fig. 5d). Overall results demonstrate that the AkaSuke/DkumLuc1 reaction is a potential orthogonal pair in combination with the AkaLumine/Akaluc reaction, becoming a good template for further refinement of dual NIR-BLI.

## Discussion

In this study, we presented the NIR luciferin analogue, AkaSuke, that is effectively catalyzed by Fluc, achieving to extend capabilities of NIR-BLI. In addition to AkaSuke, we synthesized the analogues **1b** and **1c** (Fig. 1a). A major challenge in the synthesis of bioluminescence substrate is the lack of reliable predictive methodologies for determining the structure of luciferins and catalytic properties. To assess the luminescence capacity of the luciferin analogues, we performed DFT and TD-DFT calculations of the oxy-forms (oxy-**1a–c**) of AkaSuke (**1a**) and its analogues **1b** and **1c** (Supplementary Fig. 1 and Supplementary Table 1). While these calculations indicated a potential of oxy-**1a–c** as light emitters, they alone were insufficient for comprehensive prediction of bioluminescence properties. In fact, we could not observe any luminescence signal for bioluminescence assays with **1c**. The HOMO level of oxy-**1c** (–6.11 eV) was found to be lower than those of oxy-AkaSuke (oxy-**1a**) and oxy-**1b**, while the electron density distributions of HOMO and LUMO of oxy-**1c** were similar to those of oxy-AkaSuke (oxy-**1a**) and oxy-**1b**. The non-bioluminescence behavior of **1c** is similar to that of a pyrazine-substituted luciferin analogue reported previously ^10^. That is, the oxy-form of the pyrazine-substituted luciferin analogue has a low HOMO level at –5.91 eV. The decrease in the HOMO level indicates a decrease in the electron-donating property of the π-electronic system. One of the possible explanations on their non-bioluminescence behavior is that the reduced electron-donating property of the π-electronic system induces a reduced efficiency to generate the singlet excited state of oxy-form produced from the dioxetanone intermediate by the charge transfer induced luminescence (CTIL) mechanism ^24^. Similarly, accurate prediction of bioluminescence wavelength remains a challenging. The bioluminescence wavelengths of AkaSuke and its positional isomer **1b** were 680 and 625 nm, respectively, revealing a notable shift of 55 nm. The HOMO-LUMO gaps [Δ*E*(H-L)] of oxy-AkaSuke and oxy-**1b** were 2.77 and 2.94 eV, respectively (Supplementary Table 1), confirming the correlation between the measured bioluminescence wavelengths and the calculated Δ*E*(H-L) values. Unfortunately, however, the Δ*E*(H-L) value of seMpai (Fig. 1a)was 2.96 eV ^10^, which was nearly identical to that of oxy-**1b**, yet the bioluminescence wavelength of seMpai (680 nm) was considerably red-shifted compared to that of **1b** (625 nm). These findings imply that the polar environment within the enzyme active site exerts a significant influence on the energy level of the excited state of the luminescent oxy-form. Thus, further refinements on the correlation studies that connect bioluminescence wavelengths, HOMO-LUMO gaps and substrate-enzyme interactions would be desired to provide improved prediction of bioluminescence properties of synthetic luciferins.

The *K*_m_ value of AkaSuke was considerably higher than that of AkaLumine, yet similar to that of D-luciferin in terms of kinetic constant. The high water solubility of AkaSuke may reduce its affinity for the hydrophobic active site of the enzymes. The *V*_max_ values for AkaSuke and D-luciferin were nearly identical. In this context, the *V*_max_ of the enzyme reaction is indicative of the bioluminescence quantum yield (*Φ*_BL_) and rate constant (*k*_cat_) of the luminescence reaction. Given that the luminescence quantum yield is inversely related to wavelength, the luminescence quantum yield of AkaSuke is anticipated to be lower than that of D-luciferin. Consequently, the *k*_cat_ of AkaSuke is presumed to be larger than that of D-luciferin. Moreover, the property that both *K*_m_ and *V*_max_ values are elevated suggests a direct proportionality between substrate concentration and luminescence intensity. This is supported by the data that demonstrates a strong linearity (Fig. 2a). Therefore, substantial *K*_m_ and *V*_max_ values would be one of the essential parameters for practical applications.

We demonstrated that AkaSuke improves the detection sensitivity of both cancer cells and hematopoietic cells in various deep tissues including the lung, lymph node and bone marrow of mice (Fig. 3a, b, d and e). Notably, a combination with AkaSuke and DkumLuc1 further enhanced lung tissue imaging demonstrated by comparable performance with AkaLumine/Akaluc (Fig. 4c and d), suggesting that AkaSuke/DkumLuc1 holds one of the most sensitive BLI imaging modality. However, despite generating 10 time more photons in the AkaSuke/Fluc reaction compared to AkaLumine/Fluc reaction, AkaSuke does not enhance deep brain imaging performance over AkaLumine (Fig. 2a and Supplementary Fig. 7). These results indicate poor permeability of AkaSuke to BBB as compared to AkaLumine. Thus, AkaLumine/Akaluc would be more suitable imaging system for brain as well as improved BLI tools based on marine luciferases such as the cephalofurimazine/Antares reaction ^25^. As well as brain imaging, the improved animal imaging was unsuccessful in the liver (Supplementary Fig. 8c and d). This is perhaps related to the mechanisms underlying liver autoluminescence of AkaSuke (Supplementary Fig. 8a). AAV-mediated gene expression in the liver is dominantly observed in hepatocytes ^26,27^. AkaSuke might undergo unbeneficial bioavailability in hepatocytes, as demonstrated by modest photon emission from the liver with AAV-mediated Fluc expression, but not from Fluc-expressing cancer cells in the liver (Supplementary Fig. 8c-f). Even though the detail mechanism of the liver autoluminescence is still unknown, the autoluminescence generated by AkaSuke was significantly lower than the one by AkaLumine, being suitable for high-sensitive animal imaging (Supplementary Fig. 8b). Modest autoluminescence of AkaSuke could be explained by its biodistribution if the autoluminescence mechanism is shared between AkaSuke and AkaLumine. AkaSuke is probably excluded through the kidney because of its high water solubility, whereas the high hydrophobicity of AkaLumine would lead to its accumulation in the liver where diverse metabolic enzymes might catalyze luminescence reaction of these substrates. The further mechanism study of the autoluminescence is meaningful both for improved NIR-BLI with high signal-to-noise ratio and for endogenous enzyme imaging in mammals.

We assessed dual BLI with the potential orthogonal pair of NIR synthetic bioluminescence systems (Fig. 5). In classic line, several combinations with firefly luciferases and marine luciferases were attempted to visualize two or three different targets in mice ^28–30^. Recent efforts have further pushed the strategy with marine luciferases by the development of coelenterazine analogues for NanoLuc. The dual BLI using a combination of hydrofurimazine/NanoLuc and AkaLumine/Akaluc is one of the most sensitive dual BLI system ^31^. Whereas, dual NIR-BLI systems have been developed elegantly through exploring bioluminescence systems with a firefly luciferin analogue, Infra-luciferin, displays different wavelength shift in reaction with Fluc_green and Fluc_red, demonstrating that this system enables to track cancer cells and T cell dynamics in mice ^15^. In addition, firefly luciferin analogues NH_2_-NpLH_2_ successfully construct dual NIR-BLI in combination with click beetle luciferase CBR_2pot_ ^32^. Although these systems enable simultaneous detection of dual targets through spectral separation of NIR signals, one limitation is modest photon output ^15,32^. The AkaSuke/DkumLuc1 and AkaLumine/Akaluc reactions harness rich photon outputs and robust NIR wavelength for highly sensitive in vivo imaging, offering a framework for the development of next-generation dual NIR-BLI systems by enhancing their orthogonality and improving spectral unmixing techniques.

## Methods

### Ethical Statements

Ethical approval of this study protocol for recombinant DNA experiments and animal experiments was obtained from Jichi Medical University, RIKEN and University of Miyazaki. The maximal tumor size permitted by our ethics committees or institutional review boards was 10 mm at the largest diameter in mice and was not exceeded in our experiments.

### Synthesis of AkaSuke and its analogues

AkaSuke and its analogues **1b** and **1c** were synthesized following the procedure detailed in the supporting information (Supplementary Fig. 2).

### Measurement of bioluminescence emission spectra

Bioluminescence emission spectra of D-luciferin (Promega, Madison, WI, USA), AkaLumine, seMpai, AkaSuke, **1b** and **1c** were measured using an ATTO AB-1850 spectrophotometer (ATTO Co. Ltd., Tokyo, Japan). A reaction mixture was prepared by mixing 5 μL of a substrate (100 μM), 5 μL of Fluc (QuantiLum^®^ Recombinant Luciferase, Promega, E1701) solution (1 mg/mL), and 5 μL of potassium phosphate buffer (KPB, 500 mM, pH 8.0). For the measurement of bioluminescence reactions using DkumLuc1, DkumLuc1 was expressed with Histag in E.Coli and purified to prepare DkumLuc1 solution (1 mg/mL) in 50 mM Tris-HCl containing 10% glycerol, pH 8.0. Luminescence reactions were then initiated by injecting 10 μL of ATP-Mg (200 μM) into the reaction mixture. The substrates were dissolved in 50 mM KPB (pH 6.0), Fluc solution was diluted to 1 mg/mL by 50 mM KPB containing 35% glycerol, pH 6.0, and Mg-ATP was dissolved in ultrapure water. Bioluminescence emission spectra were measured in 0.25 nm increments from 400 nm to 780 nm using 3 minutes of integration time. Light intensity was determined as the intensity of the emission spectrum at the *λ* value.

### Measurement of the kinetics analysis

For the kinetics analysis, the light emission from bioluminescence reaction was monitored with the ATTO AB-2280 luminometer for 30 seconds with sampling intervals of 0.1 seconds. Emission intensities were expressed as counts per second (cps). The *K*_m_ and *V*_max_ values of the substrates were determined from the integrated values of emission intensities and calculated using GraphPad Prism 9 (GraphPad Software, San Diego, CA, USA). For the bioluminescence reaction with Fluc, the reaction was initiated by injecting 40 μL of Mg-ATP (200 μM) into a mixture of 20 μL of a substrate solution, 20 μL Fluc solution (0.1 mg/mL), and 20 μL KPB (500 mM, pH 8.0) at room temperature. The substrates were dissolved in 50 mM KPB (pH 6.0), Fluc solution (1 mg/mL in 50 mM Tris-HCl containing 10% glycerol, pH 8.0) was diluted to 0.1 mg/mL by 50 mM KPB (pH 6.0) containing 35% glycerol, and Mg-ATP was dissolved in ultrapure water. The final concentrations of the substrates were varied from 0.02 μM to 2 mM for AkaSuke, and from 0.02 to 200 μM for **1b**, respectively.

### Cell culture

The human embryonic kidney cells HEK293T (632180, CloneTech, Mountain View, CA, USA), human pancreatic cancer cell SUIT-2 (JCRB1094, JCRB cell bank, Osaka, Japan) and human cervical cancer cells HeLa (CCL-2, ATCC, Manassas, VA, USA) were cultured in Dulbecco’s Modified Eagle’s medium (DMEM) with high glucose (Thermo Fisher Scientific, Waltham, MA, USA). The murine breast cancer cells E0771 (94A001, CH3 Biosystems, Buffalo, NY, USA) were cultured in Roswell Park Memorial Institute-1640 (RPMI) medium with L-Glutamine (FUJIFILM Wako Pure Chemical Corporation, Osaka, Japan). All media contained 10% fetal bovine serum (FBS) (Thermo Fisher Scientific), 100 IU/mL penicillin/streptomycin (Nacalai Tesque) and were used for culturing cells in a humidified incubator at 37 °C, with 5% CO_2_. All cell lines were regularly tested for mycoplasma contamination and were authenticated by morphology check and growth curve analysis.

### Plasmid construction

To construct the CSII-CMV-MCS backbone plasmid (RIKEN BRC, Saitama, Japan) for producing lentiviral particles carrying Venus-Fluc reporter gene, we amplified Venus-Fluc (luc2) cDNA and inserted it into the Not1 site of the CSII-CMV-MCS vector by using In-Fusion HD Cloning Kit (Takara Bio, Shiga, Japan). For the transient expression of luc2, Akaluc and DkumLuc1, we amplified cDNAs of luciferases and inserted into the EcoRⅤ site of pcDNA3.1(+)/myc-His (Thermo Fisher Scientific, Waltham, MA, USA). A to obtain pcDNA3.1/CMV-luc2, pcDNA3.1/CMV-Akaluc and pcDNA3.1/CMV-DkumLuc1.

### Gene transduction in cells

For stable gene transduction, we prepared lentivirus particles. The CSII plasmids were co-transfected with the packaging plasmid psPAX2 (#12260, addgene, Watertown, MA, USA), the VSV-G- and Rev-expressing plasmids (pCMV-VSV-G-RSV-Rev) (RIKEN BRC) into HEK293T cells (CloneTech) by PEI MAX (Polysciences, Warrington, PA, USA) as previously described ^26^. E0771, SUIT-2 and HeLa cells were cultured for 48 hours in the medium with the lentivirus particles and 10 µg/µL polybrene (Sigma-Aldrich). The successfully transduced cells were selected by fluorescence-activated cell sorting for expression of Venus or mCherry. For transient gene transduction, PEI MAX and plasmids (pcDNA3.1/CMV-luc2, pcDNA3.1/CMV-Akaluc and pcDNA3.1/CMV-DkumLuc1) were incubated in serum-free medium for 15 minutes and added to HEK29T cells cultured in a 6-well plate. We used the cells for cellular bioluminescence assay 24 hours after gene transduction.

### BLI of living cells

The substrates were reacted with E0771/Venus-Fluc or HEK293T transiently expressing luciferases (2 × 10^5^ cells per well) in a black 96-well plate. Bioluminescence was measured 30 seconds after adding the substrates using IVIS LuminaⅡ (PerkinElmer, Shelton, CT, USA). The following conditions were used for image acquisition: open for total bioluminescence or Cy5.5 (680 ± 10 nm) of an emission filter for NIR bioluminescence, exposure time = 10 seconds, binning = small: 4, field of view = 12.9 × 12.9 cm, and f/stop = 1. For the bioluminescence kinetics measurement in living cells, E0771/Venus-Fluc (5 × 10^4^ cells per well) was treated with the substrates in a black 96-well plate. Bioluminescence was measured at every 1 minute after adding the substrates using IVIS LuminaⅡ (PerkinElmer). The following conditions were used for image acquisition: open for total bioluminescence, exposure time = 10 seconds, binning = small: 8, field of view = 12.9 × 12.9 cm, and f/stop = 1. The all bioluminescence images were analyzed by Living Image 4.3 software (PerkinElmer) specialized for IVIS system.

### Mice

C57B/6 albino mice (female), NOD-SCID mice (male) or C.B-17 SCID mice (male) were obtained from Charles River Laboratory Japan (Yokohama, Japan). The *ff*luc mice were regularly bred with C57B/6 mice to establish a cohort for experimental use. The genotype of the offspring was verified through both fluorescence and bioluminescence detection of the cpVenus-Fluc reporter gene. Lck-Akaluc mice were described in the previous study ^22^. Lck–Cre male mice (RBRC04738, B6.Cg-Tg(Lck-cre)1Jtak) were crossed with Cre-dependent female Venus/Akaluc reporter mice (RBRC10858, C57BL/6J-Gt(ROSA)26Sorem14(CAG-Venus/Akaluc)Rbrc/#87) to generate Lck-Akaluc mice. For ROSA CAG-luc2 KI mice, a targeting vector containing a luc2 cDNA under the control of the CAG promoter into the ROSA 26 locus was constructed as described previously ^22^ and injected into the pronuclei of fertilized eggs of C57BL/6NJcl mice in combination with a ribonucleotide protein cocktail for CRISPR/Cas9 genome editing as described in the previous study ^33^. Founder mice with correct knockin of the targeting vector at the ROSA 26 locus and lacking plasmid backbone cointegration were selected to establish a ROSA CAG-luc2 KI mouse line (RBRC11209, RIKEN BRC). The ROSA CAG-luc2 KI mice were further crossed with B6.Cg-c/c Hr^hr^ mice (RBRC05798, RIKEN BRC) to introduce the albino and hairless phenotypes for imaging of the freely behaving mice with the digital color camera. The Lck-Akaluc and ROSA CAG-luc2 KI mice (male and female) were prepared to be 14-32 weeks at the experiment, and other mice used were littermates or age-matched (6-8 weeks of age) females or males. All mice were provided access to food and water *ad libitum*, and were housed in the animal facilities at Jichi Medical University and University of Miyazaki.

### BLI of mice

Bioluminescence images of lung metastasis and hematopoiesis were acquired with IVIS Spectrum at 15 minutes post-intraperitoneal injection of the indicated amount of D-luciferin, or at 3 minutes following injection of AkaSuke, unless otherwise noted. For obtaining bioluminescence emission kinetics in mice, bioluminescence images of lung metastasis were sequentially acquired with IVIS Spectrum every 3 minutes for 30 minutes after intraperitoneal injection. To compare bioluminescence production across different substrates using identical mice, images of lung metastasis were acquired by i.p. injection of AkaSuke (3 μmol/body) 6 hours after i.p. injection of D-luciferin (10 μmol/body). The following conditions were used for image acquisition: open emission filter, exposure time = 60 seconds, binning = small: 4, field of view = 12.9 × 12.9 cm, and f/stop = 1. For bioluminescence imaging of HEK293T cells in the lung, the substrates were injected intraperitoneally 15 minutes after intravenous injection of HEK293T cells (2 × 10^5^ cells / body). Then, time-course BLI were performed with IVIS Spectrum by acquiring bioluminescence imaging every 3 minutes for 15 minutes to compare the peak intensity of the different bioluminescence reactions. Peak bioluminescence intensity was typically observed at 12 minutes in the AkaSuke/DkumLuc1 imaging and at 3 minutes in the AkaLumine/Akaluc imaging. The following conditions were used for image acquisition: open emission filter, exposure time = 30 seconds, binning = medium: 8, field of view = 22.6 × 22.6 cm, and f/stop = 1. The bioluminescence images were analyzed by Living Image 4.3 software (PerkinElmer) specialized for IVIS system.

For dual NIR-BLI, Lck-Akaluc mice hair was removed before imaging. Image acquisition was accomplished by using a Lumazone system equipped with an iKon L camera (Andor Technology Ltd., UK) and a lens (YMV2595N, φ50 mm, f: 0.95, Yakumo Optical Corp.) under the control of Andor Solis (Andor Technology Ltd., UK). The bioluminescence images for detecting xenogeneic transplantation were acquired by i.p. injection of AkaSuke (3 μmol/body) 6 hours after i.p. injection of AkaLumine (1.5 μmol/body). The following conditions were used for image acquisition: exposure time = 10 seconds (for AkaSuke/DkumLuc1 imaging sessions at day1-3, 30 seconds for all other imaging sessions), Binning = 4: Readout Rate = 50 kHz. The bioluminescence images were analyzed by MetaMorph (Molecular Devices LLC., Sunnyvale, CA, USA) and quantified photon counts per 10 seconds to draw the graphs.

### Cancer metastasis models

For lung metastasis model, E0771/Venus-Fluc (5 × 10^5^ cells per 100 μL) suspended in PBS was injected from tail vein of C57B/6 albino mice (male). The experiments were performed 10-20 days after intravenous injection. This metastasis model is well-established and metastatic formation is stable. Therefore, 5 mice are adequate sample size for evaluation of bioluminescence emission from metastatic sites.

For liver metastasis model, SUIT-2/Venus-Fluc (3 × 10^5^ cells per 100 μL) suspended in PBS was injected from spleen of C.B-17 SCID mice (male). The experiments were performed 20-30 days after cancer cell injection. This metastasis model is well-established and metastatic formation is stable. Therefore, 4 mice are adequate sample size for quantitative evaluation of bioluminescence emission from metastatic sites.

### Hematopoiesis expansion model

The femurs of *ff*luc mice were harvested by removing the muscles and residual tissues surrounding the bone. We then cut the femurs at both ends and flush the bone marrow component by injecting 500 μL PBS at the one end of the bone using 29G syringe. The bone marrow component was filtered through the 35 μm cell trainer (Falcon) and centrifuged at 4 °C. The cell pellet was then rinsed with distilled water to lyse red blood cells and resuspended in PBS. For transplantation, the bone marrow cells (1 × 10^6^ cells per 100 μL) suspended in PBS was injected from tail vein of NOD-SCID mice (female). Preliminary experiments determined that the sample size of 6 mice is adequate for quantitative evaluation of bioluminescence emission from LN and BM.

### Xenogeneic transplantation model

HeLa/mCherry-DkumLuc1 (1 × 10^6^ cells per 10 μL) suspended in PBS was mixed with an equal volume of Geltrex (Invitrogen) and subcutaneously injected into Lck-Akaluc mice (male and female). This transplantation model is well-established and immune rejection response is stable. Therefore, 4 mice are adequate sample size for quantitative evaluation of bioluminescence emission from xenotransplantation sites.

### Statistics and Reproducibility

Data are presented as means ± standard error of the mean (SEM) and were statistically analyzed with two-tailed Student’s t-test or one-way ANOVA followed by Bonferroni’s multiple comparison. P-values < 0.05 were considered statistically significant. Statistical analyses were performed using GraphPad Prism 9 (GraphPad Software, San Diego, CA, USA).

## Acknowledgements

We thank Kurogane Kasei Co., Ltd., for providing AkaLumine, AkaLumine-HCl (TokeOni) and seMpai. We thank Y. Suzuki (Jichi Medical University, Japan) for technical assistance of plasmid construction and animal experiment. We thank K. Kobata and R. Kawasaki (University of Miyazaki, Japan) for technical assistance of cell culture and animal care. We thank Center for Instrumental Analysis, Tokyo University of Pharmacy and Life Sciences for HRMS analysis. We thank University of Electro-Communications Coordinated Center for UEC Research Facilities, supported by “Advanced Research Infrastructure for Materials and Nanotechnology in Japan (ARIM)” of the Ministry of Education, Culture, Sports, Science and Technology (MEXT) in proposal Number JPMXP1222UE5277, JPMXP1223UE5277. This work was supported by Japan Society for the Promotion of Science (JSPS) KAKENHI Grant Numbers JP20K22724 to R. S.-M, JP20H03178 to S.I., JP15H05948 to S. A. M., and the CREST program of the Japan Science and Technology Agency Grant Number 214660001 to T.K.

## Author contributions

R.S.-M., N.K., R.O., R.I., H.Ab., H.Ao. and S.A.M. designed and synthesized luciferin analogues. R.S.-M., S.I., N.K., N.Y., G.K., R.O., T.H., H.Ao. and S.A.M. characterized luciferin-luciferase reactions in test tubes. R.S.-M., N.K., N.Y., G.K. and T.H. calculated oxy-luciferin analogues. T.K., A.C.-E, T.I. characterized luciferin-luciferase reactions in living cells. S.I., T.K., Y.Y, K.Y. and T.N. performed animal experiments. K.O. and T.Su. cloned DkumLuc1. Y.K., M.T., H.H. and T.O. provided AAV tools. Y.M. performed flowcytometry analysis. R.S.-M., S.I., T.K., N.K., D.S., T.Sa., S.S., H.N., T.O., T.N., T.H., H.Ao. N.T., A.M., S.A.M. discussed data. R.S.-M, S.I., T.K., Y.K., T.O., T.N., T.H., N.T., A.M., S.A.M. wrote the paper. T.K. and S.A.M supervised the project. All authors reviewed the paper draft.

## Data Availability

The raw data will be available within two weeks upon reasonable request to T. Kuchimaru and S. Maki. Source data are also provided with this paper.

