## Supplementary Figs and Tables for "A bright synthetic near-infrared luciferin enhances the capabilities of deep-tissue bioluminescence imaging using firefly luciferases"

Saito-Moriya et al.

- -Supplementary Figures
- -Supplementary Tables

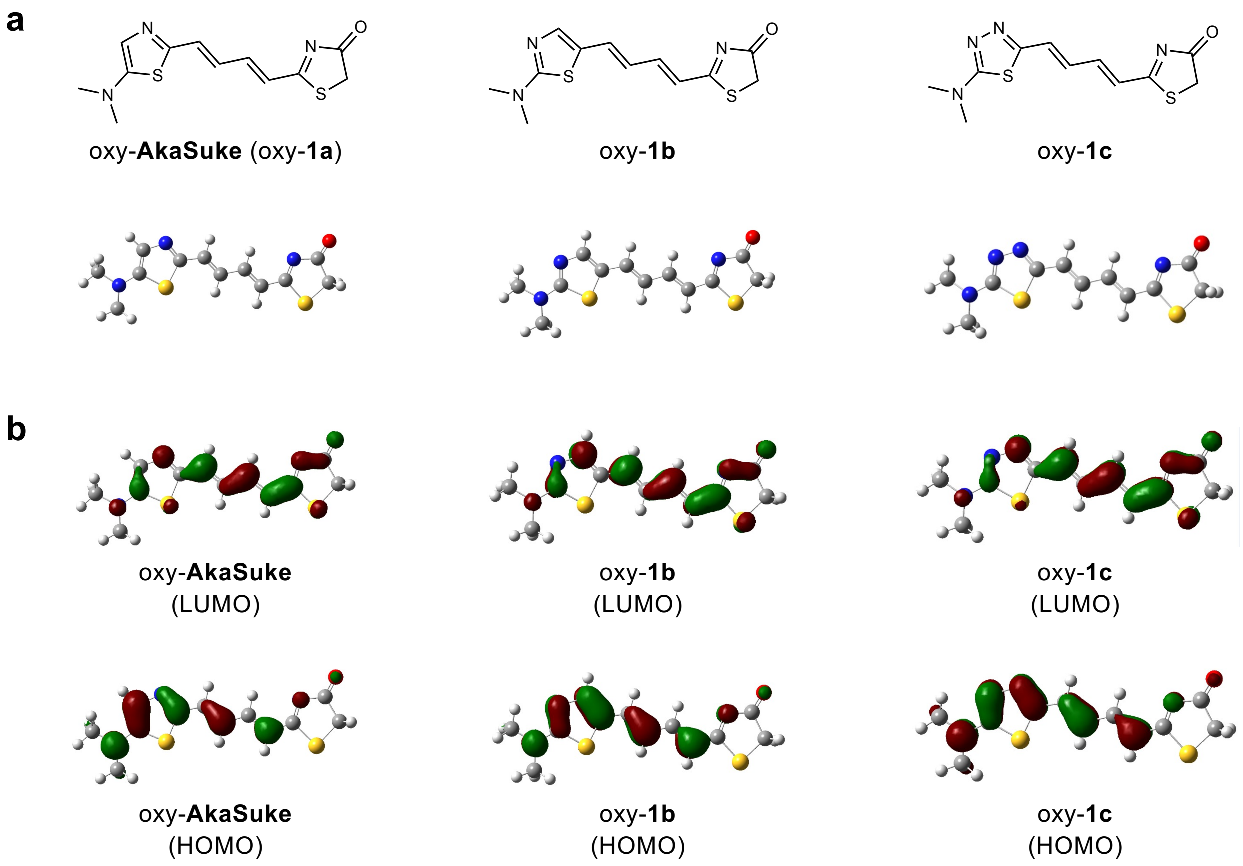


**Supplementary Fig. 1 Optimized structures of oxy-AkaSuke, oxy-1b and oxy-1c (a) and their HOMOs and LUMOs (b)**.

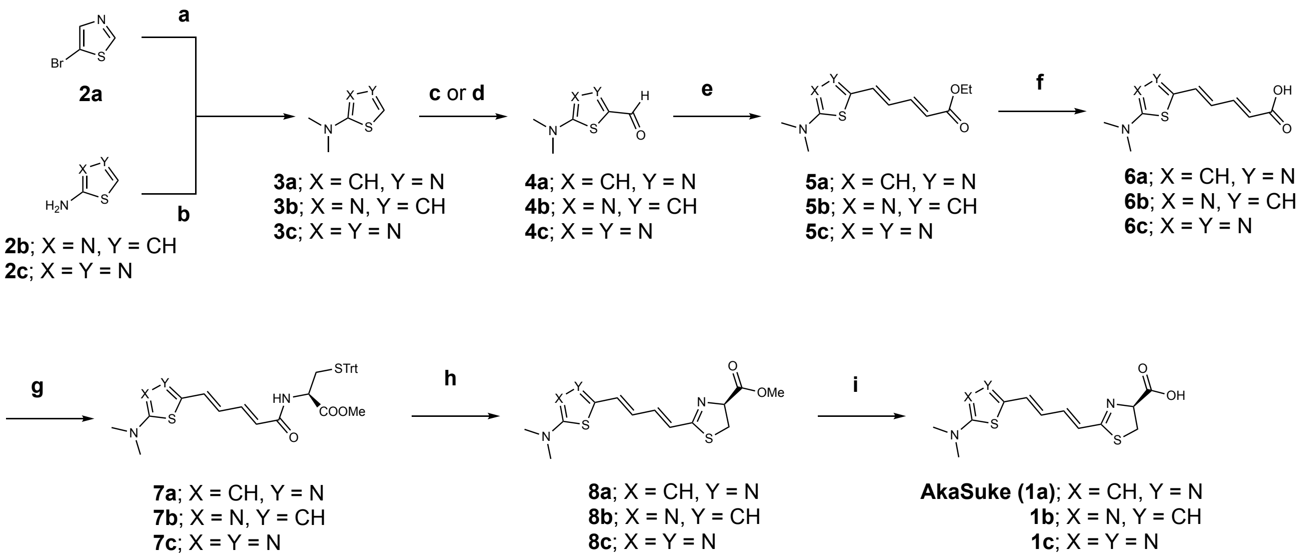


**Supplementary Fig. 2 Schematic of the synthesis route of firefly luciferin analogues AkaSuke (1a), 1b and 1c.** A detailed synthetic protocol is provided in the Supplementary Methods section.

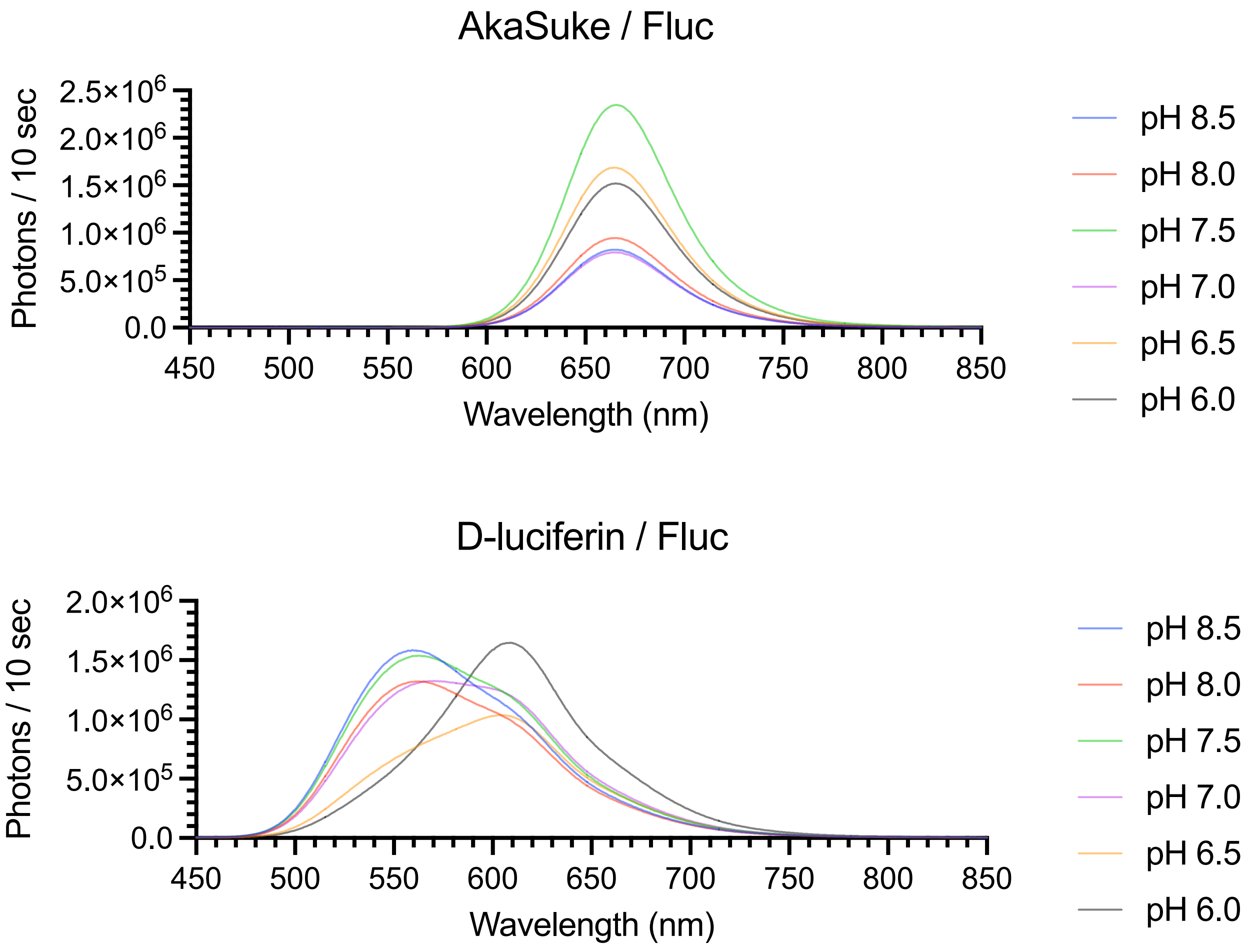


**Supplementary Fig. 3 Bioluminescence spectrum of AkaSuke/Fluc and D-luciferin/Fluc reactions under the varied pH conditions.** Source data are provided as a Source Data file.

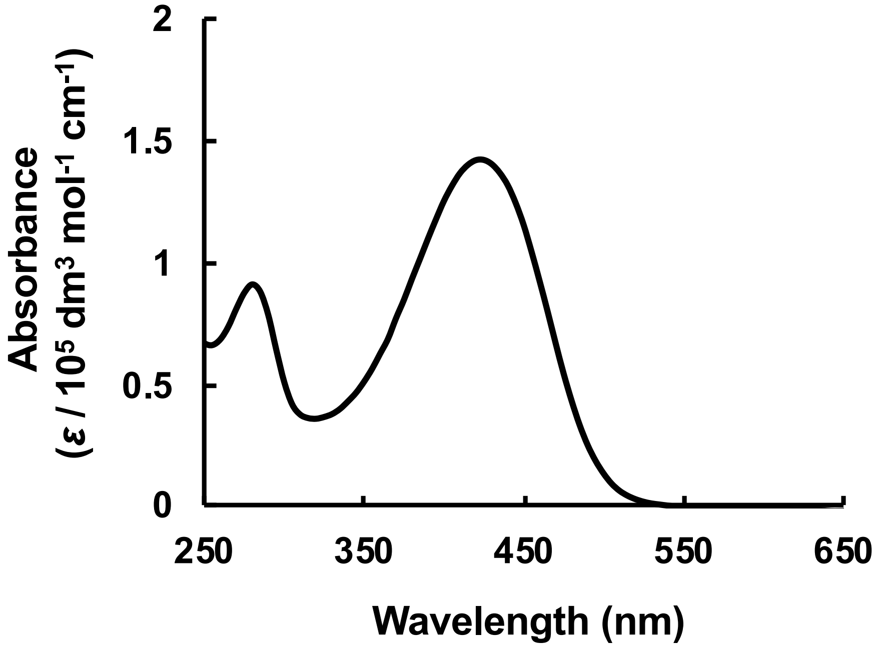


**Supplementary Fig. 4 UV-vis absorption spectrum of AkaSuke in PBS (pH = 7.4).**
Source data are provided as a Source Data file.

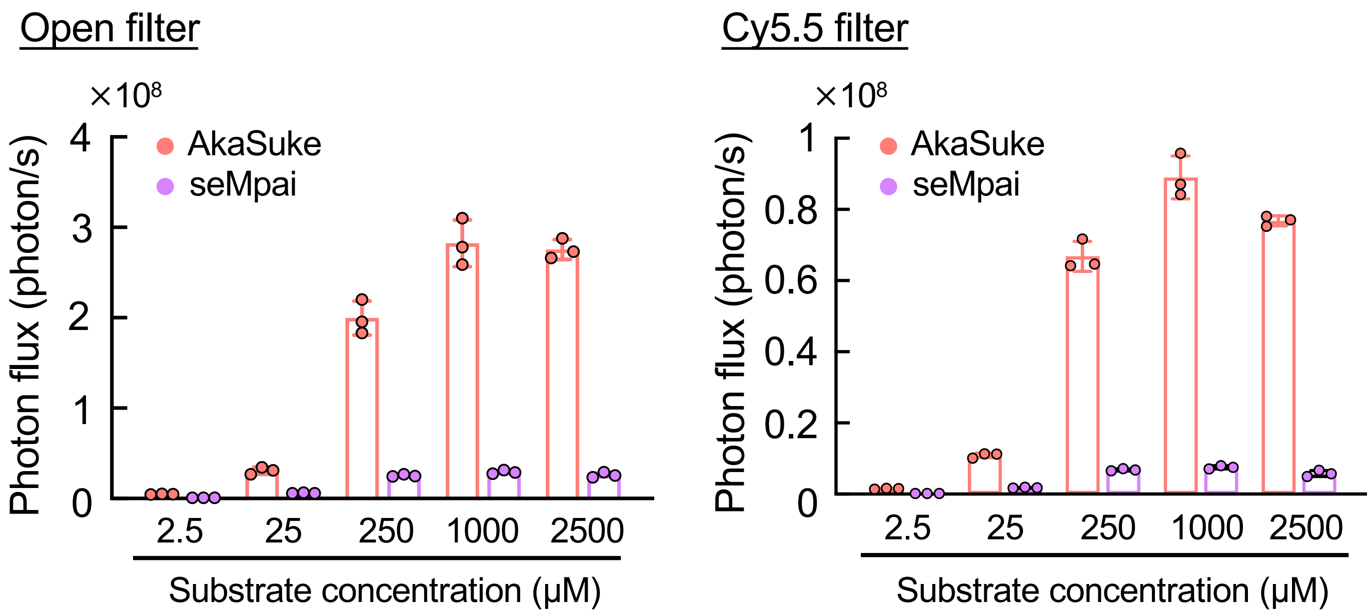


**Supplementary Fig.5 Bioluminescence reactions of AkaSuke/Fluc and seMpai/Fluc in living cells**. Bioluminescence emission from living cells expressing Fluc (HEK293T/Fluc) in reaction with the substrate at indicated concentrations. Bioluminescence emissions were measured using an open or Cy5.5 filter. Data were statistically analyzed with two-tailed Student's t-test (n = 3 biologically independent samples). Data are presented as mean values ± SEM. Source data are provided as a Source Data file. All experiments were repeated at least twice, and similar results were reproducible.

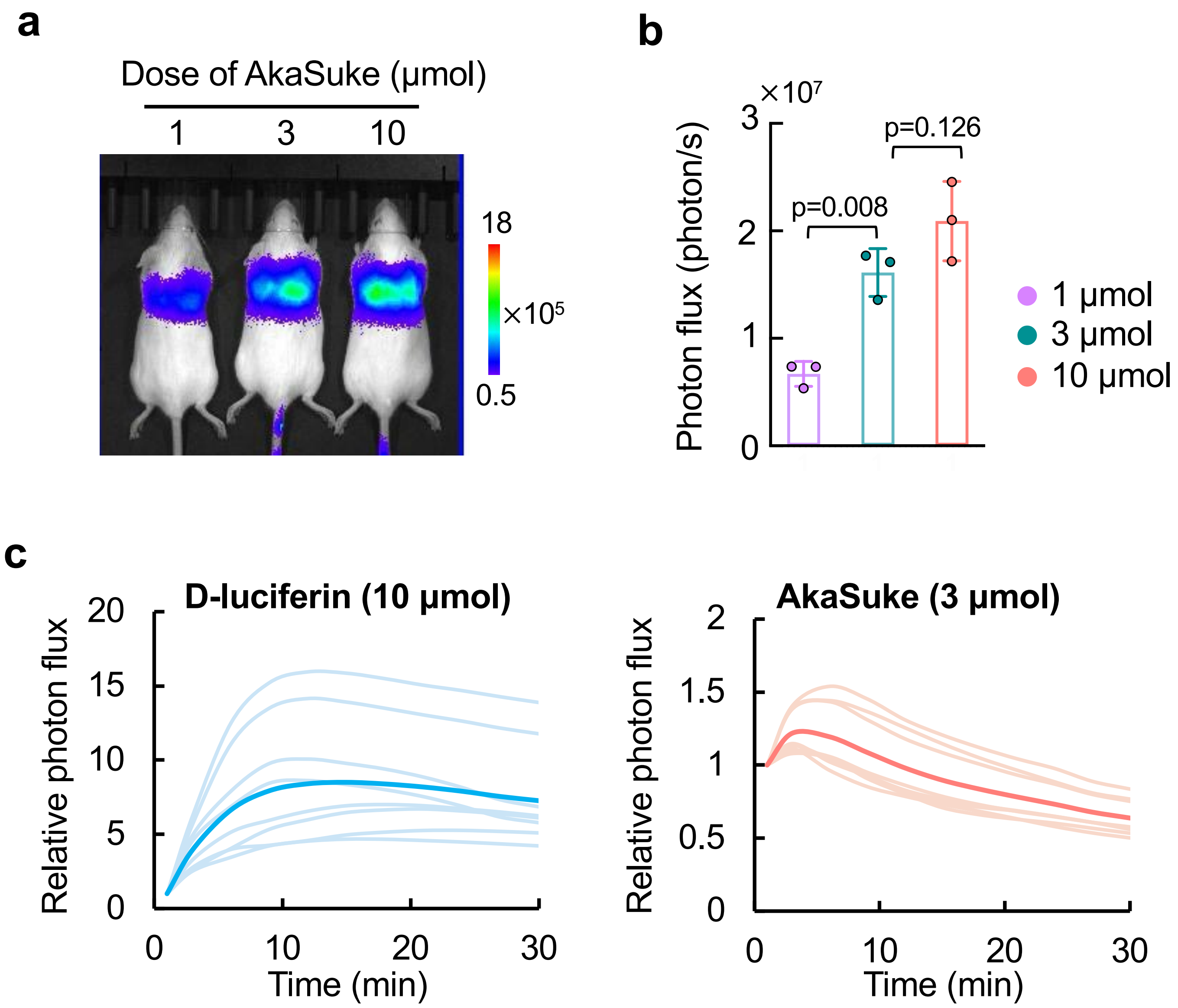


**Supplementary Fig. 6 NIR-BLI of living mice using AkaSuke. a** Representative bioluminescence images of mice intravenously injected with E0771/Venus-Fluc cells. The images were acquired by i.p. injection of varied dose of AkaSuke. **b** Quantitative analysis of bioluminescence emission from the lung in mice injected with varied dose of AkaSuke (n = 3 independent mice, p-value for 1 μmol vs 3 μmol / 3 μmol vs 10 μmol, one-way ANOVA followed by Bonferroni's multiple comparison). Source data are provided as a Source Data file. **c** Time-course changes of bioluminescence emission from E0771/Venus-Fluc lung metastasis after i.p. injection of D-luciferin (10 μmol/body) or AkaSuke (3 μmol/body) (n = 8 independent metastatic lesions in 5 mice). Thick curve represents the mean value, which is calculated from thin curves in each graph. Source data are provided as a Source Data file. All experiments were repeated at least twice, and similar results were reproducible.

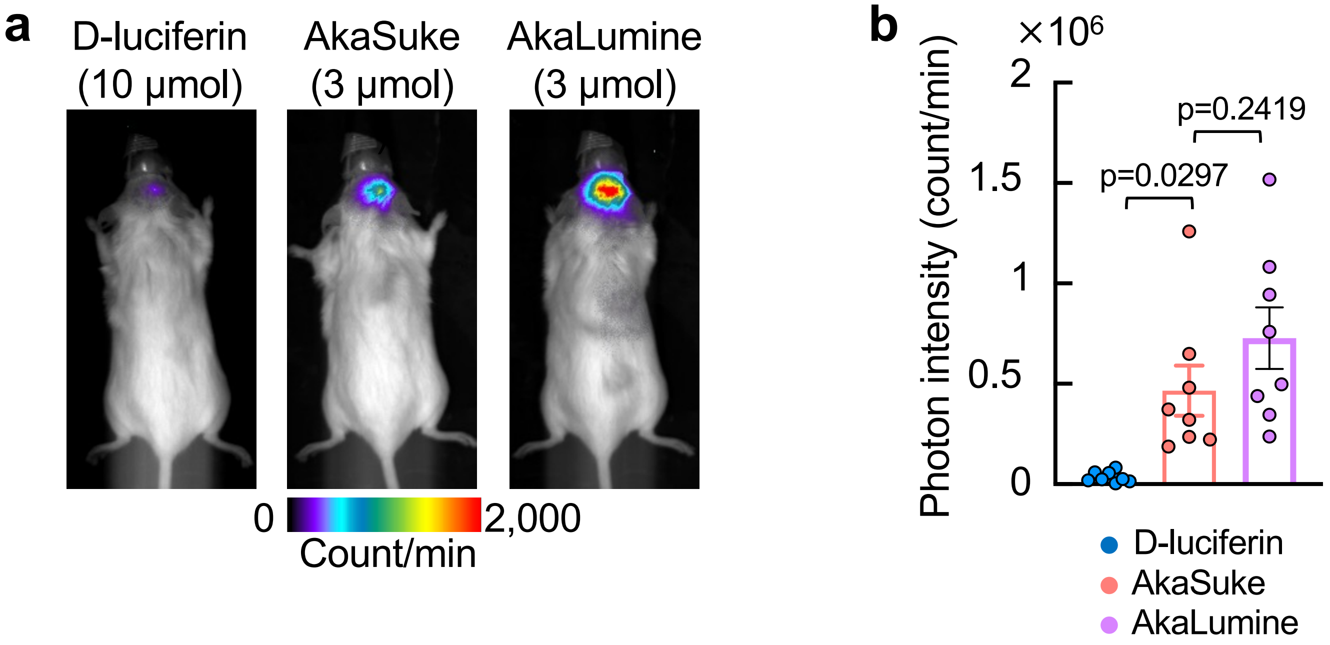


**Supplementary Fig. 7 NIR-BLI of the deep brain of living mice using AkaSuke. a** Representative bioluminescence images of mice locally injected with AAV encoding Fluc at striatum. The images were acquired by i.p. injection of the substrate at indicated dose. **b** Quantitative analysis of bioluminescence emission from the brain in mice injected with each subtrate (n = 8 independent mice, p-value for D-luciferin vs AkaSuke / AkaSuke vs AkaLumine, one-way ANOVA followed by Bonferroni's multiple comparison). Source data are provided as a Source Data file. Source data are provided as a Source Data file. All experiments were repeated at least twice, and similar results were reproducible.

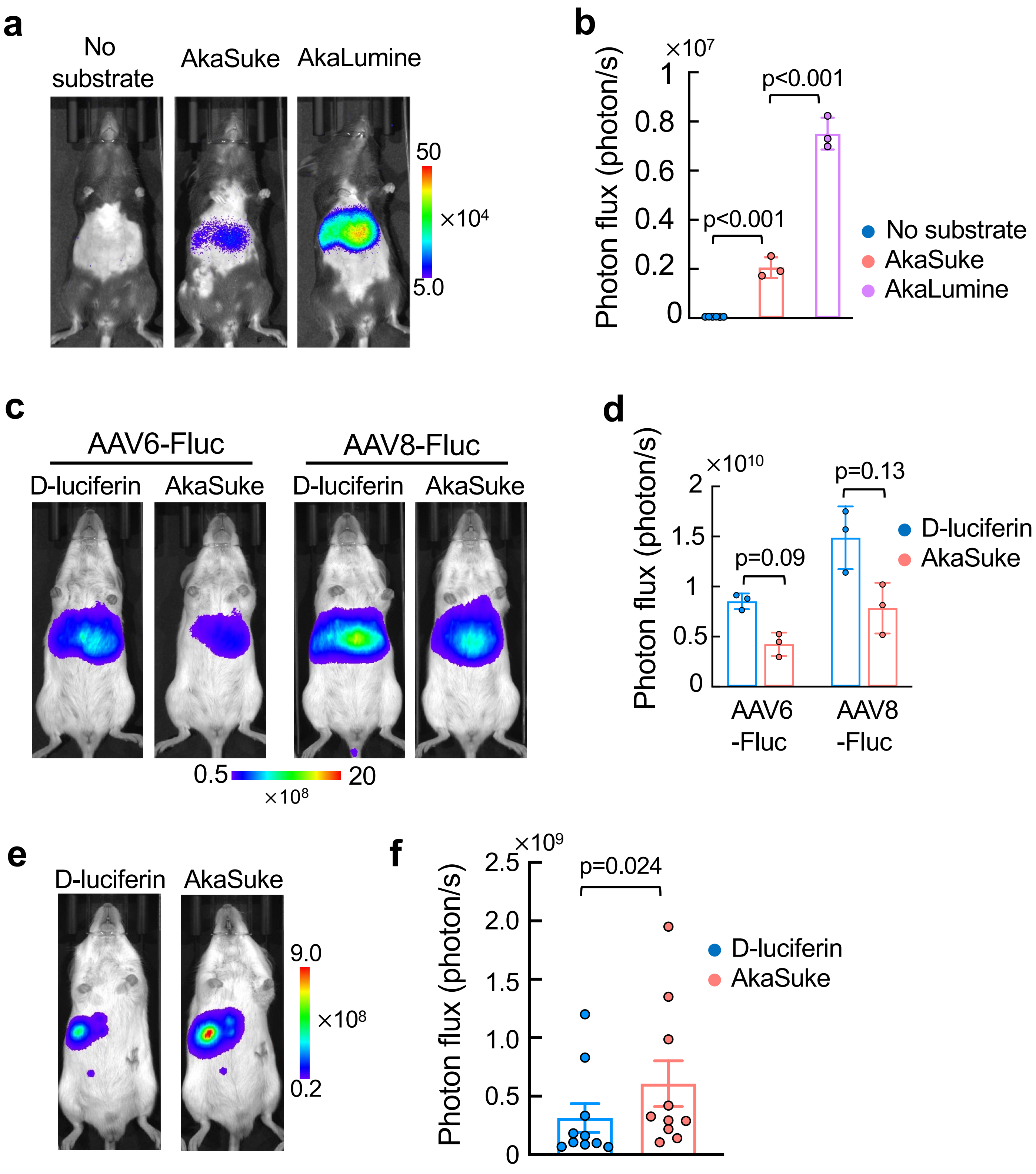

a
No substrate
AkaSuke
AkaLumine
50
×10^4
5.0
b
×10^7
Photon flux (photon/s)
1
0.8
0.6
0.4
0.2
0
p<0.001
p<0.001
No substrate
AkaSuke
AkaLumine
c
AAV6-Fluc
AAV8-Fluc
D-luciferin
AkaSuke
D-luciferin
AkaSuke
0.5
20
×10^8
d
×10^10
Photon flux (photon/s)
2
1.5
1
0.5
0
p=0.09
p=0.13
D-luciferin
AkaSuke
AAV6 -Fluc
AAV8 -Fluc
e
D-luciferin
AkaSuke
9.0
×10^8
0.2
f
×10^9
Photon flux (photon/s)
2.5
2.0
1.5
1.0
0.5
0
p=0.024
D-luciferin
AkaSuke

**Supplementary Fig. 8 Bioluminescence emission of AkaSuke in the liver. a** Representative bioluminescence images of autoluminescence emission from the liver of mice without Fluc. Mice were injected with 3 μmol of AkaSuke or 1.5 μmol of AkaLumine. **b** Quantitative analysis of autoluminescence emission in the liver (n = 6 for no substrate, n = 3 for AkaSuke/AkaLumine, p-value for no substrate vs AkaSuke and AkaSuke vs AkaLumine, one-way ANOVA followed by Bonferroni's multiple comparison). Source data are provided as a Source Data file. **c** Representative bioluminescence images of mice transduced with AAVs carrying Fluc. The images were acquired by i.p. injection of D-luciferin (10 μmol/body) and AkaSuke (3 μmol/body). **d** Quantitative analysis of bioluminescence emission in the liver of mice transduced with AAVs carrying Fluc (n = 3, p-value for no D-luciferin vs AkaSuke, two-tailed Student's *t* test). Source data are provided as a Source Data file. **e** Representative bioluminescence images of mice harboring liver metastasis (SUIT-2/Venus-Fluc). The images were acquired by i.p. injection of D-luciferin (10 μmol/body) and AkaSuke (3 μmol/body). **f** Quantitative analysis of bioluminescence emission from liver metastasis generated with D-luciferin and AkaSuke (n = 10 independent metastatic lesions in 4 mice, p-value for D-luciferin vs AkaSuke, two-tailed Student's *t* test). Source data are provided as a Source Data file. All experiments were repeated at least twice, and similar results were reproducible.

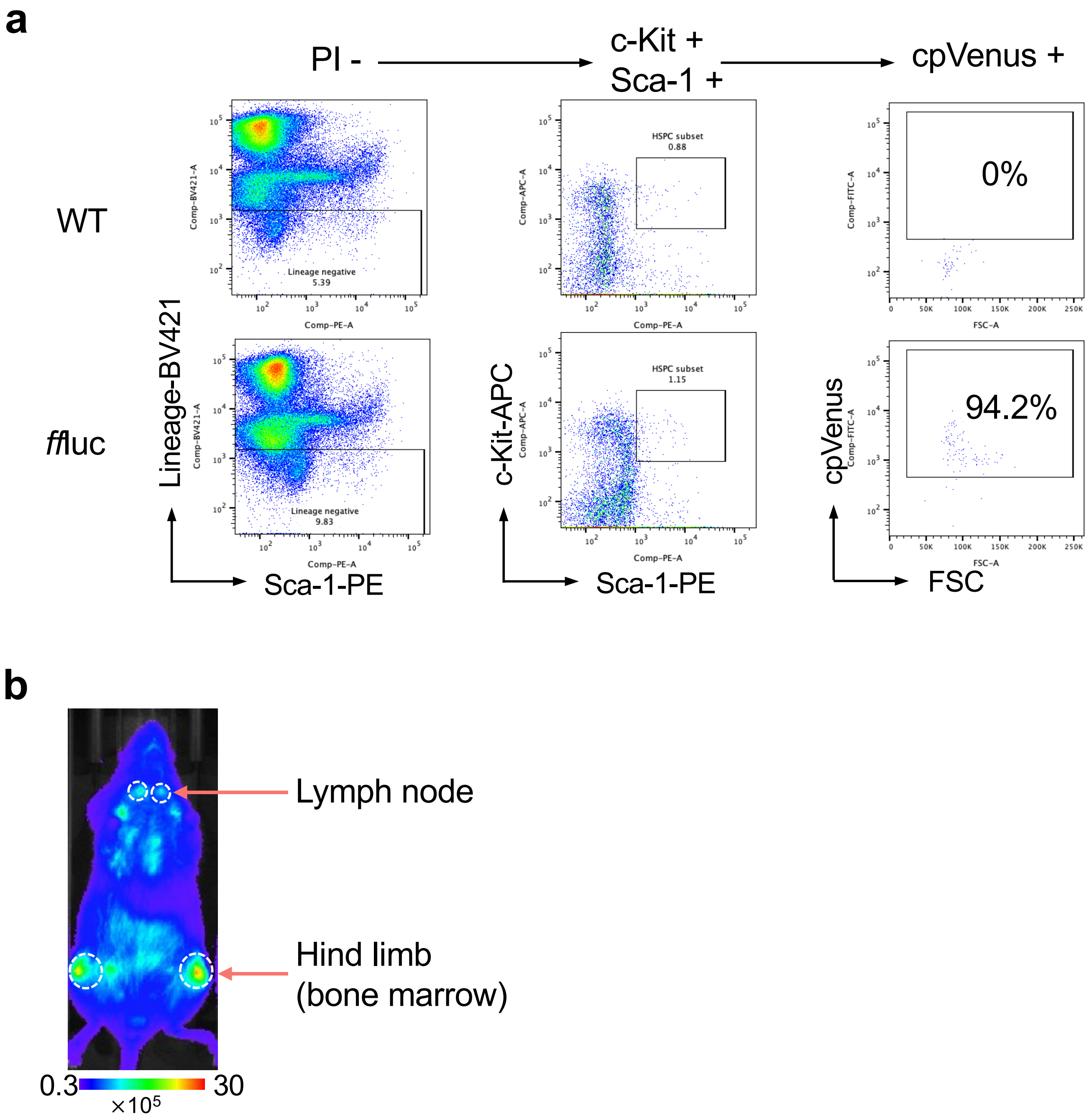


**Supplementary Fig. 9 Transplantation of hematopoietic stem cells isolated from *ff*luc Tg mice. a** A representative FACS profile of mouse bone marrow is displayed. Among living cells (propidium iodide-negative cells), the Sca-1-positive and c-Kit-positive fraction in the lineage marker (CD3e, CD45R, Gr-1, CD11b, and Ter119)-negative fraction was defined as hematopoietic stem/progenitor cells (HSPCs) or the KSL fraction. Regarding KSL cells, we compared the expression of Venus in cells derived from WT (Top) and *ff*luc-Venus mice (Bottom). The experiment was repeated twice, and similar results were reproducible. **b** A representative bioluminescence image indicates anatomical region of interest (white-dashed circles) for the quantitative analysis in Fig.3e.

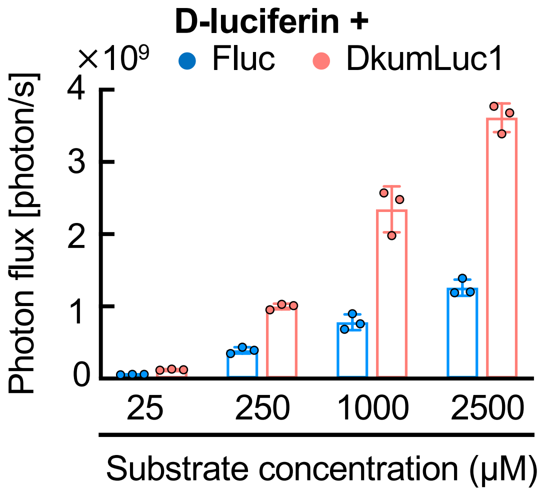


**Supplementary Fig. 10 Bioluminescence activity of DkumLuc1 in living cells.** Bioluminescence emission from living HEK293T cells expressing Fluc or DkumLuc1 treated with D-luciferin at indicated concentrations. Source data are provided as a Source Data file. The experiment was repeated twice, and similar results were reproducible.

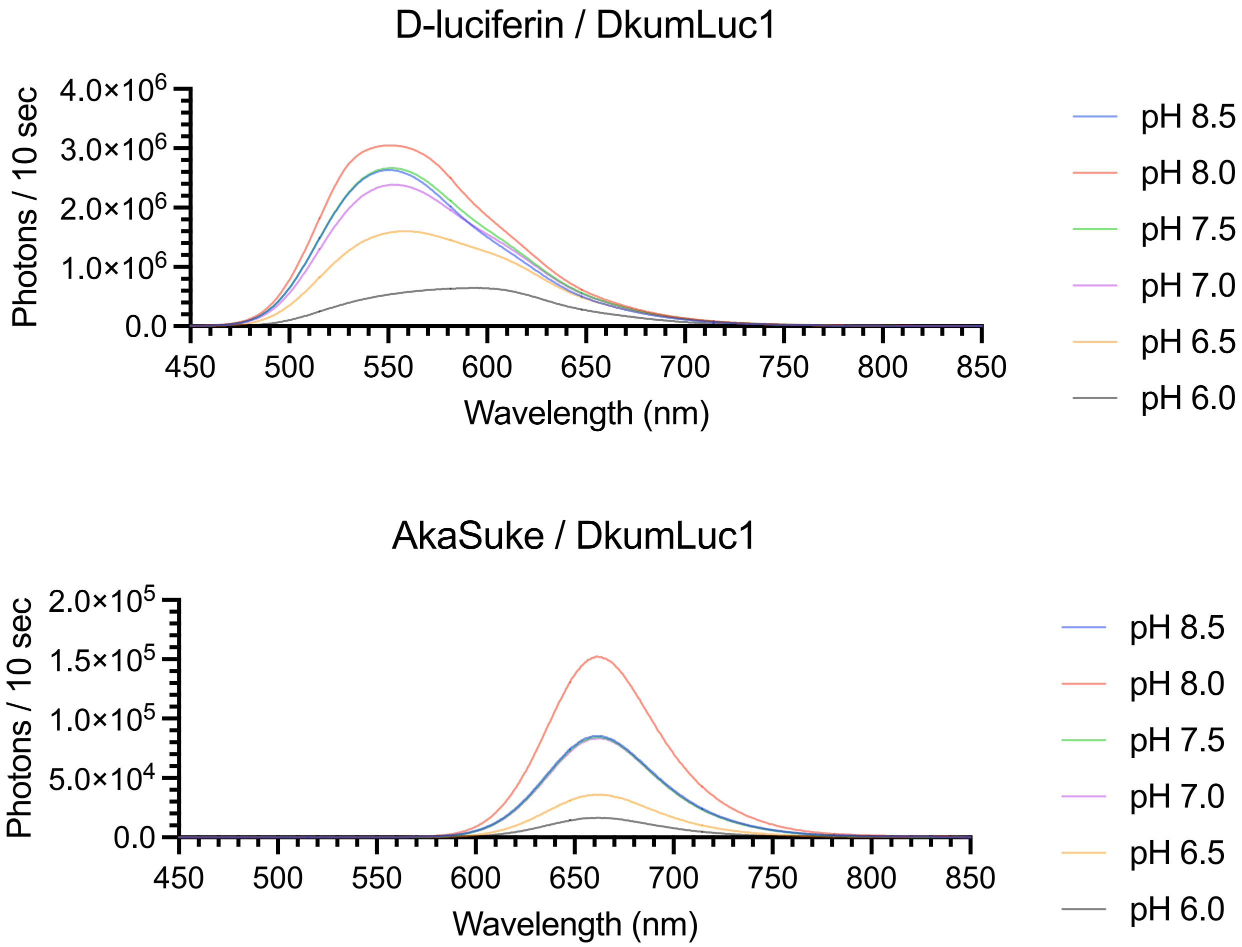


**Supplementary Fig. 11 Bioluminescence spectrum of AkaSuke/DkumLuc1 and D-luciferin/DkumLuc1 reactions under the varied pH conditions.** Source data are provided as a Source Data file.

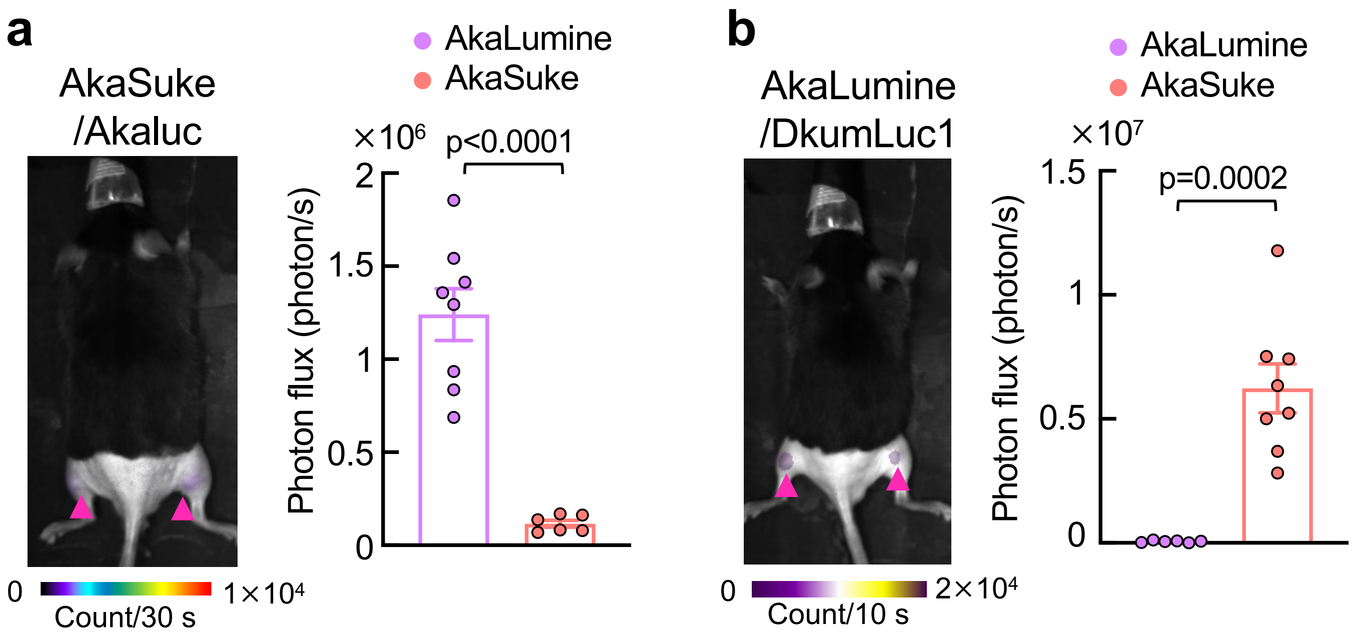


**Supplementary Fig. 12 The reactivity of AkaSuke/Akaluc and AkaLumine/DkumLuc1 in BLI of mice. a** AkaSuke was intraperitoneally injected (3 μmol/body) into Lck-Akaluc mice 7 days after subcutaneous injection of naive human cells (HeLa) at hind limbs. Quantitative analysis of bioluminescence emission from the xenotransplantation sites indicated by magenta arrow-heads (n = 6 independent xenotransplantation sites in 3 mice for the AkaSuke/Akaluc reaction) was performed with bioluminescence signals generated by the AkaLumine/Akaluc reaction at day7 in Fig. 4e (n = 8 independent xenotransplantation sites in 4 mice). The p-value was calculated for AkaLumine vs AkaSuke by two-tailed Student's *t* test. Source data are provided as a Source Data file. **b** AkaLumine was intraperitoneally injected (1.5 μmol/body) into wild-type mice 2 days after subcutaneous injection of HeLa/DkumLuc1 at hind limbs. Quantitative analysis of bioluminescence emission from the xenotransplantation sites indicated by magenta arrow-heads (n = 6 independent xenotransplantation sites in 3 mice for the AkaLumine/DkumLuc1 reaction) was performed with bioluminescence signals generated by the AkaSuke/DkumLuc1 reaction at day2 in Fig. 4e (n = 8 independent xenotransplantation sites in 4 mice). The p-value was calculated for AkaLumine vs AkaSuke by two-tailed Student's *t* test. Source data are provided as a Source Data file. All experiments were repeated at least twice, and similar results were reproducible.

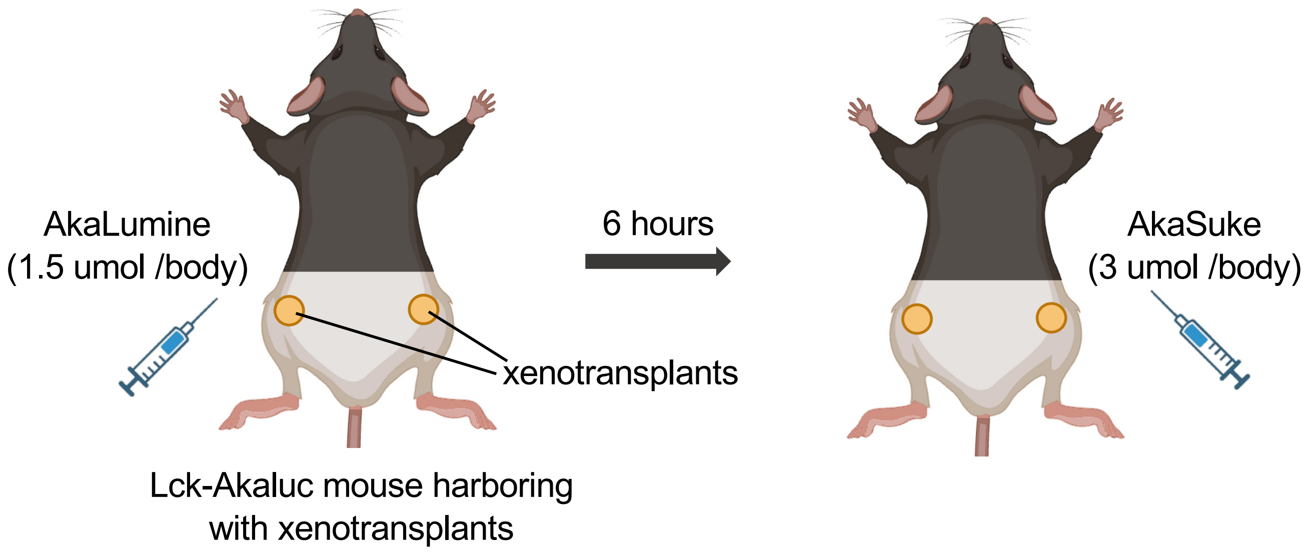


**Supplementary Fig. 13 A schematic diagram of dual target detection by animal NIR-BLI.** The hair on the lower body of Lck-Akaluc mice was removed before imaging. The bioluminescence images for detecting xenogeneic transplantation were acquired by i.p. injection of AkaSuke (3 μmol/body) 6 hours after i.p. injection of AkaLumine (1.5 μmol/body).

**Supplementary Table1 DFT and TD-DFT calculation data for oxy-AkaSuke, oxy-1b and oxy-1c.**

| Compound | HOMO (eV) | LUMO (eV) | Δ*E*(H-L)[a] (eV) | $\lambda_{tr}$ (nm) (*f*)[b] | Configuration[c] |
|---|---|---|---|---|---|
| oxy-**AkaSuke** (oxy-**1a**) | -5.66 | -2.89 | 2.77 | 452 (1.29) | H→L (0.71) |
| oxy-**1b** | -5.69 | -2.74 | 2.94 | 424 (1.43) | H→L (0.70) |
| oxy-**1c** | -6.11 | -3.07 | 3.05 | 419 (1.20) | H→L (0.70) |

[a]Energy difference between HOMO and LUMO. [b]Wavelength estimated from the energy of the $S_0 \rightarrow S_1$ transition. Oscillator strength (*f*) is in parenthesis. [c]Configuration of excitation. Coefficients are in the parentheses. H and L denote HOMO and LUMO, respectively.

**Supplementary Table 2 Heats of formation of oxy-AkaSuke, oxy-1b and oxy-1c with DFT using B3LYP/6-31+G(d).**

| Compound | Heat of formation (hartree) |
|---|---|
| oxy-**AkaSuke** (oxy-**1a**) | -941853.2003 |
| oxy**-1b** | -941860.7900 |
| oxy**-1c** | -951910.9236 |

**Supplementary Table 3 Cartesian Coordinates (in Å) of oxy-AkaSuke.**

| | | x | y | z |
|---|---|---|---|---|
| 1 | C | -0.2507281 | 4.5773649 | 0.2334134 |
| 2 | C | 0.7527884 | 5.4753898 | -0.1294915 |
| 3 | S | 2.1096484 | 4.5783103 | -0.7846271 |
| 4 | H | -1.1954591 | 4.854436 | 0.6829344 |
| 5 | N | 0.0078023 | 3.2672663 | -0.0047857 |
| 6 | C | 1.1940206 | 3.0678403 | -0.5359558 |
| 7 | C | 1.7046076 | 1.7707967 | -0.8687698 |
| 8 | H | 1.0136089 | 0.9628617 | -0.6331303 |
| 9 | C | 2.9158433 | 1.4771533 | -1.4243371 |
| 10 | H | 3.6137056 | 2.2787303 | -1.6680161 |
| 11 | C | 3.3249351 | 0.1355712 | -1.7106822 |
| 12 | H | 2.6320698 | -0.6698134 | -1.4712268 |
| 13 | C | 4.5251064 | -0.2019831 | -2.2643049 |
| 14 | H | 5.2488891 | 0.5691404 | -2.5201643 |
| 15 | C | 4.8950085 | -1.570103 | -2.5364176 |
| 16 | S | 6.5043231 | -1.8652199 | -3.2679887 |
| 17 | N | 4.1428769 | -2.6041446 | -2.288184 |
| 18 | C | 4.7363083 | -3.8125209 | -2.6460511 |
| 19 | O | 4.2288039 | -4.9091486 | -2.5104491 |
| 20 | N | 0.7399471 | 6.8437722 | -0.0607402 |
| 21 | C | 1.9984394 | 7.5795209 | -0.1044264 |
| 22 | H | 1.7830591 | 8.6418614 | -0.2433223 |
| 23 | H | 2.5900983 | 7.4538982 | 0.8165879 |
| 24 | H | 2.6009785 | 7.2455317 | -0.9548914 |
| 25 | C | -0.3725411 | 7.4659293 | 0.64878 |
| 26 | H | -0.3446706 | 7.2614876 | 1.7310402 |
| 27 | H | -0.3299324 | 8.5462152 | 0.4922343 |
| 28 | H | -1.3226163 | 7.1012818 | 0.2466027 |
| 29 | C | 6.1528422 | -3.6498284 | -3.2562637 |
| 30 | H | 6.1675384 | -4.0608412 | -4.2692486 |
| 31 | H | 6.8815054 | -4.1907249 | -2.6465117 |

**Supplementary Table 4 Cartesian Coordinates (in Å) of oxy-1b.**

| | | x | y | z |
|---|---|---|---|---|
| 1 | C | 0.7406327 | 5.4522418 | -0.1842285 |
| 2 | S | 2.2063747 | 4.4569186 | -0.3032233 |
| 3 | C | 1.1739496 | 3.0217485 | -0.3843068 |
| 4 | C | 1.6499728 | 1.6815693 | -0.5079373 |
| 5 | H | 0.8638865 | 0.9257951 | -0.5330682 |
| 6 | C | 2.9478453 | 1.2607916 | -0.5948483 |
| 7 | H | 3.7575148 | 1.9903456 | -0.574003 |
| 8 | C | 3.2985784 | -0.1200332 | -0.7151607 |
| 9 | H | 2.4896646 | -0.8493862 | -0.736489 |
| 10 | C | 4.5759558 | -0.5934475 | -0.8052917 |
| 11 | H | 5.4198592 | 0.093115 | -0.788853 |
| 12 | C | 4.8702414 | -1.9992093 | -0.9246574 |
| 13 | S | 6.5937211 | -2.4816527 | -1.0354362 |
| 14 | N | 3.9741506 | -2.94544 | -0.9578743 |
| 15 | C | 4.5233765 | -4.21838 | -1.0781725 |
| 16 | O | 3.8878256 | -5.2544839 | -1.1294121 |
| 17 | N | 0.8075269 | 6.8062694 | -0.1221307 |
| 18 | C | 2.0864503 | 7.4757694 | 0.0651981 |
| 19 | H | 1.9650894 | 8.5360635 | -0.1693898 |
| 20 | H | 2.4558852 | 7.382848 | 1.0978842 |
| 21 | H | 2.8378412 | 7.0637589 | -0.6165158 |
| 22 | C | -0.4076254 | 7.5595146 | 0.1858755 |
| 23 | H | -0.5296126 | 7.6942012 | 1.2705822 |
| 24 | H | -0.3444536 | 8.5419645 | -0.2912349 |
| 25 | H | -1.2723849 | 7.0200406 | -0.1992309 |
| 26 | C | 6.0730165 | -4.2209349 | -1.1450467 |
| 27 | H | 6.4007523 | -4.6720583 | -2.0854801 |
| 28 | H | 6.4769064 | -4.8064229 | -0.3147342 |
| 29 | N | -0.3891362 | 4.7669268 | -0.1828329 |
| 30 | C | -0.1429793 | 3.4358698 | -0.3009326 |
| 31 | H | -0.9835439 | 2.7488176 | -0.3225141 |

**Supplementary Table 5 Cartesian Coordinates (in Å) of oxy-1c.**

| | | x | y | z |
|---|---|---|---|---|
| 1 | C | -0.0664095 | 5.2834102 | 0.1989648 |
| 2 | S | -1.0293455 | 3.8149572 | 0.3842488 |
| 3 | C | 0.2669494 | 2.9968376 | -0.5142668 |
| 4 | C | 0.2874777 | 1.5968549 | -0.8409257 |
| 5 | H | 1.1678739 | 1.2971791 | -1.4067727 |
| 6 | C | -0.6557576 | 0.6716408 | -0.5150783 |
| 7 | H | -1.5376224 | 0.9674936 | 0.0533723 |
| 8 | C | -0.5445552 | -0.7093634 | -0.8913069 |
| 9 | H | 0.3332362 | -1.0143606 | -1.4589329 |
| 10 | C | -1.4679616 | -1.6607693 | -0.5829744 |
| 11 | H | -2.3599438 | -1.3995515 | -0.0171035 |
| 12 | C | -1.3206985 | -3.0439808 | -0.9810134 |
| 13 | S | -2.6144846 | -4.1856919 | -0.5044494 |
| 14 | N | -0.3144897 | -3.5116692 | -1.6597061 |
| 15 | C | -0.415057 | -4.8803363 | -1.9133641 |
| 16 | O | 0.4012169 | -5.5381383 | -2.527205 |
| 17 | N | -0.4295631 | 6.4776799 | 0.7305834 |
| 18 | C | -1.7789412 | 6.6784752 | 1.2404827 |
| 19 | H | -1.792465 | 7.5868052 | 1.8479864 |
| 20 | H | -2.5175484 | 6.7844003 | 0.4310701 |
| 21 | H | -2.0740448 | 5.8406996 | 1.8803179 |
| 22 | C | 0.342124 | 7.6651008 | 0.3588465 |
| 23 | H | -0.0096477 | 8.0906121 | -0.5920668 |
| 24 | H | 0.2334791 | 8.4125659 | 1.1496746 |
| 25 | H | 1.3923496 | 7.3942431 | 0.2510852 |
| 26 | C | -1.6989735 | -5.5239792 | -1.3291626 |
| 27 | H | -1.4279193 | -6.3103562 | -0.6196442 |
| 28 | H | -2.2902434 | -5.9660543 | -2.1355425 |
| 29 | N | 1.0535679 | 5.096943 | -0.485344 |
| 30 | N | 1.230934 | 3.8164485 | -0.8655205 |

**Supplementary Table 6 $K_m$, $K_i$ and $V_{max}$ of AkaSuke/DkumLuc1 and AkaLumine/Akaluc.**

| | AkaLumine | AkaSuke |
|---|---|---|
| **AkaLuc** | $K_m$ = 2.11 μM | $K_m$ = 3.24 μM |
| | $V_{max}$ = 6.42 × $10^7$ | $V_{max}$ = 1.82 × $10^6$ |
| **DkumLuc1** | $K_m$ = 0.557 μM | $K_m$ = 8.21 μM |
| | $V_{max}$ = 2.96 × $10^5$ | $V_{max}$ = 1.67 × $10^7$ |

| Enzyme<br>Substrate<br>Inhibitor | **DkumLuc1**<br>**AkaSuke**<br>**AkaLumine** | **Akaluc**<br>**AkaLumine**<br>**AkaSuke** |
|---|---|---|
| **$K_m$ (μM)** | 8.74 | 2.12 |
| **$K_i$ (μM)** | 14.6 | 19.4 |
| **$V_{max}$** | 1.78 × $10^7$ | 6.39 × $10^7$ |
